# A Rescorla-Wagner Drift-Diffusion Model of Conditioning and Timing

**DOI:** 10.1101/184465

**Authors:** André Luzardo, Eduardo Alonso, Esther Mondragón

## Abstract

Computational models of classical conditioning have made significant contributions to the theoretic understanding of associative learning, yet they still struggle when the temporal aspects of conditioning are taken into account. Interval timing models have contributed a rich variety of time representations and provided accurate predictions for the timing of responses, but they usually have little to say about associative learning. In this article we present a unified model of conditioning and timing that is based on the influential Rescorla-Wagner conditioning model and the more recently developed Timing Drift-Diffusion model. We test the model by simulating 10 experimental phenomena and show that it can provide an adequate account for 8, and a partial account for the other 2. We argue that the model can account for more phenomena in the chosen set than these other similar in scope models: CSC-TD, MS-TD, Learning to Time and Modular Theory. A comparison and analysis of the mechanisms in these models is provided, with a focus on the types of time representation and associative learning rule used.

**Author Summary:** How does the time of events affect the way we learn about associations between these events? Computational models have made great contributions to our understanding of associative learning, but they usually do not perform very well when time is taken into account. Models of timing have reached high levels of accuracy in describing timed behaviour, but they usually do not have much to say about associations. A unified approach would involve combining associative learning and timing models into a single framework. This article takes just this approach. It combines the influential Rescorla-Wagner associative model with a timing model based on the Drift-Diffusion process, and shows how the resultant model can account for a number of learning and timing phenomena. The article also compares the new model to others that are similar in scope.

## 1 Introduction

Classical conditioning theories aim to understand how associations between stimuli are learned. Ever since Pavlov (1927) the process of association formation has been understood to depend crucially on the temporal relations between stimuli (Savastano and Miller, 1998; Balsam et al., 2006; Kirkpatrick, 2013). Yet, classical conditioning theories have so far struggled to work when time is taken into account as an attribute of the stimulus representation. The study of time as a mental representation is the object of a separate area of study known as interval timing. Interval timing theories have produced a rich variety of time representations (Gibbon et al., 1984; Killeen and Fetterman, 1988; Machado, 1997; Staddon and Higa, 1999; Matell and Meck, 2004), and therefore are a natural place to look for ways to integrate time into classical conditioning. In this paper we first analyse previous efforts in this direction before introducing a new hybrid classical conditioning and timing model.

The process of association formation is understood to be of fundamental survival value for both human and non-human animals. Prediction, which forms the core of classical conditioning, allows the organism to adapt to significant events in its surroundings. A prototypical experiment in classical conditioning, a type of associative learning, involves a neutral stimulus and an unconditioned stimulus (US) which is capable of eliciting an unconditioned response (UR). After repeated pairings of both stimuli in a specified order and temporal distance, the neutral stimulus comes to elicit a response similar to the UR. This response is called the conditioned response (CR) and the neutral stimulus is said to have become a conditioned stimulus (CS). Classical conditioning theories typically conceptualize this process as the formation of a link (association) between the internal representations of CS and US. Their basic building blocks are (Pearce and Bouton, 2001; Brandon et al., 2002): (a) the representations of stimuli, and (b) a learning rule to update the association weights between these representations. Although most theories do not attempt to find neurophysiological correlates, these constructs are nonetheless commonly assumed to be instantiated by (a) neural activity in the form of spike rates, and (b) synaptic plasticity (Moore, 2002; Klopf, 1988; Gallistel and Matzel, 2013). These have found some support in the neuroscientific literature, particularly studies of the role of dopamine in reward prediction (Schultz et al., 1997; Dayan and Niv, 2008; Niv, 2009; Eshel, 2016). However it is important to note that there is still no widely accepted complete neural mechanism for classical conditioning and that most theories stay at the computational level of explanation.

Stimulus representations are generally thought of as neural activation that is elicited by the stimulus, which may linger for a short time as a ‘trace’ after stimulus offset. Representations are commonly one of two types: molar or componential. Molar (or elemental) trace theories treat the stimulus as a single conceptualized unit whose activity is usually assumed to peak quite early following stimulus onset, and then gradually decrease (Hull, 1943; Wagner, 1981; Sutton and Barto, 1981; Schmajuk and Moore, 1988; McLaren and Mackintosh, 2000; Harris and Livesey, 2010). In contrast, componential trace theories break down the CS representation into smaller units, each capable of being associated with the US, with some units more active early during the CS and others late, but all leaving a trace after activation (Desmond and Moore, 1988; Grossberg and Schmajuk, 1989; Vogel et al., 2003; Ludvig et al., 2008). Learning rules may be classified according to different criteria. An important period in the recent history of the field gave rise to one of these criteria. Prior to 1970’s conditioning used to be rooted in the stimulus-response tradition, which attributed crucial importance to the temporal pairing, or contiguity, of stimuli for the development of associations. The linear operator learning rule (Hull, 1943) is one of the products of that period. In the late 1960’s and early 1970’s important experimental discoveries using compound stimuli, that is, a stimulus formed by combining other individual stimuli, showed the contiguity view to be incomplete (Rescorla, 1988; Gallistel and Gibbon, 2001). These compound experiments indicated that the formation of associations also depended on the reinforcement history of the individual elements forming the compound stimulus. This led to the development of new learning rules (Rescorla and Wagner, 1972; Mackintosh, 1975; Pearce and Hall, 1980) capable of combining individual reinforcement histories in compounds, which the linear operator rule cannot. The first, and arguably still the most influential, of these learning rules is the Rescorla-Wagner (RW, Rescorla and Wagner, 1972).

The CR is usually not a single event. Organisms time their responses so that they emerge gradually during the duration of the CS and reach maximum frequency or intensity around the time of reinforcement. Interval timing theories have attempted to provide an account for this *timing* of the CR. One of the fundamental properties of timing behaviour is that it is approximately timescale invariant, i.e. the whole response distribution scales with the interval being timed (Gibbon, 1977; Allman et al., 2014).One of the consequences of timescale invariance is that the coefficient of variation, that is the standard deviation divided by the mean, of the dependent measure of timing is approximately constant. A number of timing models have put forth explanations for timescale invariance and other timing properties (how time is encoded, how it is stored in memory and how it gets translated into behaviour) by recourse to an internal pacemaker. The most influential pacemaker-based timing theory to date is Scalar Expectancy Theory (SET, Gibbon et al., 1984; Gibbon and Church, 1984). The pacemaker is supposed to mark the passage of time by emitting pulses. These pulses can be gated to an accumulator via a switch which closes at the start of a relevant interval and opens when the interval is finished. The accumulator count is kept in working memory. At the end of the interval the current count is transferred to a long-term reference memory. Behaviour is guided by the action of a comparator which actively compares the count in working memory to the one retrieved from reference memory.

In spite of the considerable overlap, interval timing and classical conditioning are not easily integrated. Most conditioning theories are trial-based, that is they consider the trial as the unit of time. A trial is generally taken to be the state where a CS is present (or CSs in compound) and which may or may not contain a US (or USs). The most influential model in this category is the Rescorla-Wagner (RW, Rescorla and Wagner, 1972). In order to account for different stimulus durations, trial-based theories like RW must resort to some sort of time discretization, usually by subdividing the trial into ‘mini-trials’. Each mini-trial is treated as a trial in its own right, which are then used to update associative links. This gives rise to the problem of deciding on a particular discretization. Also, given that humans experience time passing as a continuous flow, it is unlikely that animals discretize their conditioning experience in such a way. A more realistic approach to timing is taken by real-time theories. These theories attempt to formalize the concept of a continuous flow of time.

The Temporal Difference model (TD, Sutton and Barto, 1990,9) was one of the earliest and still most influential real-time classical conditioning model. It may be thought of as a real-time version of RW. When used with stimulus representations such as the Complete Serial Compound (CSC, Moore et al., 1998), Microstimuli (MS, Ludvig et al., 2008,0) and the Simultaneous and Serial Configural-cue Compound (SSCC, Mondragón et al., 2014) it is capable of reproducing some timing phenomena like the gradual increase in anticipatory responding that occurs before a signalled reinforcer, and the lower response rates observed during longer CSs. However, only MS-TD has a time representation capable of approximating the most fundamental property of timing, timescale invariance. Another issue with the stimulus representations for TD is that their approach to timing resembles the strategy used by trial-based models, i.e. they all split the stimulus into a number of smaller units or states, the number of which being directly proportional to the duration of the stimulus. Given that conditioning is observed in a timescale that ranges from milliseconds to hours (Kehoe and Macrae, 2002, p. 189) this can lead to a very high number of units being required. The stimulus as a whole no doubt is a complex entity, and the brain may be employing a large number of neurons to represent it, but to dedicate so many resources only for timing might not be the most energy-efficient strategy. Also, TD and its stimulus representations do not usually account for a change in timing that is not tied to reinforcement. Animals time the occurrence of different events, such as onset and offset of stimuli (see for example Meck and Church, 1984), but TD usually only allows for the timing of rewards.

On the other hand, timing models have made even fewer attempts at integrating aspects of classical conditioning. A notable exception is the Learning to Time (LeT, Machado, 1997; Machado et al., 2009) model. It represents the passage of time by transitioning between internal states according to a stochastic pacemaker, an idea borrowed from an earlier timing model called the Behavioural Theory of Time (Killeen and Fetterman, 1988). Learning takes place by associating reinforcement presentation with the current internal state according to the linear operator, a standard classical conditioning rule. LeT offers an account of the basic dynamics of association formation, but it cannot explain cue-competition phenomena like blocking. In a blocking procedure, a CS is first paired with a US until a CR is acquired. The same CS is then presented together with a novel CS and both are paired with the US for a few trials. If the novel CS is now presented alone it elicits little or no responding, and so it is said to be blocked by the first CS. LeT’s learning rule, the linear operator, has largely been supplanted by RW in classical conditioning modelling because it cannot explain cue-competition phenomena. Like TD, LeT also employs a representation that requires as many units as time-steps, making it a resource-intense model.

Modular Theory (MoT, Guilhardi et al., 2007; Kirkpatrick, 2002) is a timing model which because of its explicit goal of integrating timing and learning may be called a hybrid theory. MoT has introduced novelties that allow it to account for some aspects of the dynamics of classical conditioning that LeT cannot. Its architecture is different than the connectionist one (states or units connected by modifiable links) assumed by RW, TD and LeT. Instead, it uses a more cognitive architecture, with separate information processing stages that deal with perception, memory and decision. It postulates two separate memories: a pattern memory which stores CS durations, and a strength memory which stores the associative strength between each pattern memory and the US. This separation allows MoT to deal with more complex situations involving the dynamics of learning during acquisition and extinction. However, MoT also relies on the linear operator to update its strength memory, which, like LeT, prevents it from accounting for cue-competition phenomena.

Although the models mentioned above, namely TD, LeT and MoT, have accomplished a great deal in terms of bringing together timing and conditioning, they each have their different strengths and weaknesses as we have touched above. In this paper we introduce a model that tries to address some of these weaknesses while preserving the strengths. More specifically, the model has the following strengths. It represents time in real-time. Like MoT and unlike LeT and TD, its time representation does not require an arbitrary large number of units or states. Similarly to TD but unlike LeT and MoT, it uses a learning rule that preserves the main features of RW which allow it to account for compound phenomena. It can time the onset and offset of all stimuli, not only of rewards, and store a memory for each. It includes two update rules: one for timing that is updated by time-markers, and another for associations that is updated by the US. Hence, simple stimulus exposure causes the model to learn and store its duration. This capability is not present in models that depend only on an associative learning rule to also learn about time, such as TD and LeT.

This new model is essentially a way to connect one of the most influential classical conditioning theories, the Rescorla-Wagner model (Rescorla and Wagner, 1972), with a recently developed timing theory called Timing Drift-Diffusion Model (TDDM, Rivest and Bengio, 2011; Simen et al., 2011). The TDDM is based on the drift-diffusion model, widely used in decision making theory, and it provides an adaptive time representation that has commonalities with pacemaker-based models like SET and LeT (Simen et al., 2013). These models postulate the existence of a pacemaker that emits pulses at a regular rate, which are then counted to mark the passage of time. To preserve timescale invariance they either postulate a specific type of noise in the memory saved for intervals and a ratio-based decision process (SET) or adapt the rate of pulses (LeT). The TDDM takes the latter route but sets a fixed threshold on pulse counting. To emphasize the unification of these two theories we call our proposal the Rescorla-Wagner Drift-Diffusion Model (RWDDM).

We evaluate RWDDM based on how well it can simulate the behaviour of animals in a number of experimental procedures. Many classical conditioning phenomena have been identified which collectively represent a significant challenge for any single model to explain. A recent list (Alonso and Schmajuk, 2012) has compiled 12 categories, which include acquisition, extinction, conditioned inhibition, stimulus competition, preexposure effects, temporal properties, among others. Of particular interest to a theory of timing and conditioning are phenomena that involve elements of both timing and conditioning. As we detail later, we have searched the literature for documented effects that can challenge the main mechanisms embodied in RWDDM.

We proceed by first introducing the new model. We compare its formalism with four models that have similar scope, namely CSC-TD, MS-TD, MoT and LeT. In the results section we present the phenomena we will simulate, followed by the results of our simulations, and compare them to the current explanations given by LeT, MoT and TD.

## 2 Model

We follow most classical conditioning theories in conceptualizing the conditioning process as the formation of an association between the internal representations of CS and US. Arguably, one of the most influential rules describing the evolution of this association through training is the Rescorla-Wagner (Rescorla and Wagner, 1972) rule. As mentioned previously, other models exist which have a similar scope to RW, both trial based (Mackintosh, 1975; Pearce and Hall, 1980) and real-time (Buhusi and Schmajuk, 1999; McLaren and Mackintosh, 2000,0). However, our goal was to take advantage of TDDM’s time representation, so we sought a theoretical associative framework that could incorporate such a representation. Since trial-based conditioning theories lack any time representation, they are a natural place to start. Out of those theories the RW is perhaps the simplest whilst also retaining the greatest possible explanatory power. Its basic formalism consists of the following rule for updating associative strength:

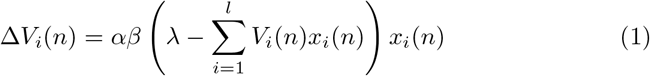

where *V_i_(n)* denotes associative strength for CS_i_ at trial *n, λ* the asymptote of learning which is set by the US representation, *x_i_* (*n*) which marks the presence (*x_i_* = 1) or absence (*x_i_* = 0) of the i-th CS representation at trial *n*, 0 < *α* < 1 a learning rate set by the CS and 0<*β*<1 a learning rate set by the US. The summation term in the equation (1) sums over all CSs present in the trial. The top panel of figure 1 shows a diagram of a basic neural net for classical conditioning which serves as the architectural framework for both RW and RWDDM. The RW rule is used to update the links *V*_1_, …, *V*_*l*_ that connect the CS input nodes CS_1_,…,CS_*l*_. The summation term in the RW rule is represented in the diagram as a summation unit or junction Σ, that sums the inputs it receives from the CSs *i = 1,…, l* present in the trial. This sum allows RW to combine (additively) the reinforcement history of each individual CS present in a compound trial. In the neural network literature, equation (1) is also referred to as the Widrow-Hoff rule (Widrow and Hoff, 1960) and the Least-Means-Square (LMS; Sutton, 1992). The relationship to the LMS rule is easier to see if we let 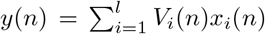 be the output of a learning unit that aims to predict a target *λ* given inputs xi by adapting the weights *V_i_*. In classical conditioning, *λ* represents the maximum learning driven by a given outcome (the US), *x_i_* is the CS and *V_i_* the associative strength. If we let *δ(n) = *λ*-y(n)* be the error between output and US, equation (1) can be obtained with the method of gradient descent by minimizing the squared error *δ*^2^(*n*) with respect to the weight *V_i_.*

**Figure 1:**
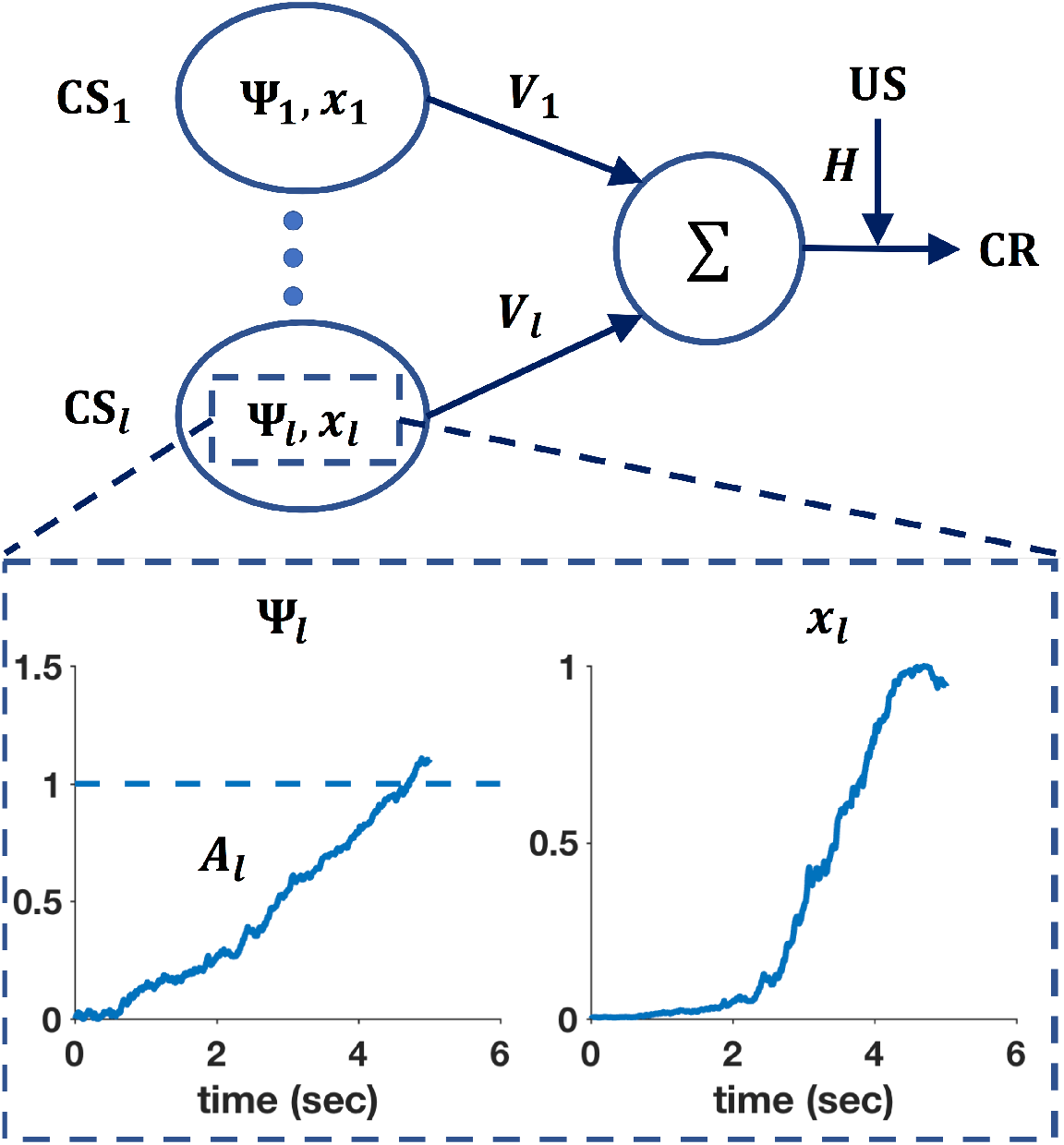
Connectionist diagram of RWDDM. Each CS unit is connected to a summing junction (labelled Σ) via a modifiable link *V*. The output of the summing junction is the CR. The US is represented as a teaching signal with a fixed weight *H*. Each CS unit has its own timer Ψ and representation *x*. The bottom panel shows a zoomed-in view of the timer Ψ_*l*_ and CS representation *x_l_* associated with CS_*l*_. The timer slope *A_l_* is tuned to a 5-second CS duration.

In spite of the relative success in explaining a wide range of conditioning phenomena (for a list of successes, and failures, see Miller et al., 1995), the Rescorla-Wagner rule lacks a mechanism to account for the microstructure of real-time responding during conditioning procedures. In terms of the order of CS-US presentation conditioning procedures may be either forward (CS followed by US) or backward (US followed by CS). Two common types of forward conditioning are delay and trace. In delay conditioning the US always occurs a fixed time after CS onset. In trace conditioning the US occurs at a fixed duration after CS offset. After sufficient training with delay or trace conditioning, responding begins some time after CS onset, increases rapidly in frequency until it reaches a maximum level where it stays until US onset (Gormezano et al., 1983). The RW rule alone does not account for CR level as a function of time. This role is usually fulfilled by the choice of CS representation. We base our choice on a timing model called Timing Drift-Diffusion Model (TDDM, Simen et al., 2011; Rivest and Bengio, 2011; Luzardo et al., 2013; Balcı and Simen, 2016). We chose the TDDM because it possesses a number of interesting features. It is part of a family of pacemaker based models like SET and LeT (Simen et al., 2013) which are arguably two of the most successful timing theories to date. The TDDM is a modified version of the drift-diffusion models that have been extremely successful at modelling reaction time in decision making tasks (Ratcliff, 1978; Voss et al., 2013). Evidence of climbing neural activity related to timing that resembles the TDDM has been extensively reported (Komura et al., 2001; Leon and Shadlen, 2003; Brody et al., 2003; Wittmann, 2013; Jazayeri and Shadlen, 2015). The TDDM consists of a drift-diffusion process with an adaptive drift or rate. The drift-diffusion process is defined by a continuous random walk called Wiener diffusion process. The two main components of Wiener diffusion are the drift and the normally distributed noise. The Wiener diffusion process may be visualized by imagining a two-dimensional grid with time in the horizontal axis and displacement on the vertical axis. If we imagine a purely linear and non-random walk that starts at the origin and moves up at a constant rate then the resulting walk would be a straight line and the drift would be equal to the slope of the line. With normally distributed noise, the walk becomes a random walk and it looks like a jagged curve, since at each time step there is now only a probability that the displacement will be up or down. For the purposes of timing, the slope is always positive and the random walk can be interpreted as a noisy accumulator (or timer) Ψ(*t*), which starts at the beginning of a salient stimulus and stops (and resets) at the end. In a conditioning experiment the CS is usually the most salient stimulus in the uneventful context of the conditioning chamber, so it is well placed to serve as a time marker. When timing starts, accumulator increments are performed at each time-step according to

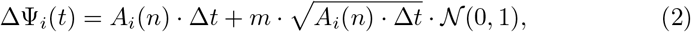

where *A_i_* (*n*) is the rate (slope) of accumulation for CS*_i_* in trial *n, m* is a noise factor, ∆*t* is the time-step size and N (0, 1) denotes a sampling from the standard normal distribution. An interval is timed by the rise in the accumulator to a certain fixed threshold, say Ψ*_i_* (*t*) = *θ*. The TDDM adjusts to new intervals by keeping the threshold fixed but adapting the rate of accumulation *A_i_ (n)*. The bottom left panel of figure 1 shows a typical trajectory (or realization) of a CS’s TDDM timer after one 5-second trial.

In its original formulation (Rivest and Bengio, 2011; Simen et al., 2011) the accumulation process was not allowed to continue beyond the threshold value *θ*, a constraint that gave rise to two distinct rules for rate adaptation, one for when the US arrived earlier than expected and another for when it arrived later. The constraint fixing a maximum level of accumulation was driven by the neurophysiological assumption that a linear neural accumulator is not likely to continue to perform effectively beyond a certain level. The neural implementation so far proposed for TDDM’s linear accumulator (Simen et al., 2011) is based on a feedback control mechanism that is tuned to balance excitation and inhibition in a neuron population. Tuning of this kind requires great computational precision, which may not be easily kept for very long in a biological system. Neurophysiology notwithstanding, we will drop that requirement here for simplicity and use instead only one update rule. We demonstrate how this single update rule can be derived by the method of gradient descent. The model learns a new interval by adapting its slope *A_i_* so that the accumulator Ψ*_i_* reaches the threshold value θ at the target time *t^∗^*, which may be the time of reinforcement for example. The target slope will therefore be *θ/t^∗^*. The error *δ*(*n*) between the target slope and the current slope is *δ*(*n*) = *θ/t^∗^–A_i_* (*n*). By minimizing the squared error *δ*^2^(*n*) using gradient descent we can derive the slope update rule. The squared error as a function of *A_i_* forms a curve. Moving in the direction opposite the slope of this curve and taking a step of size *α_t_* /2 we form the equation:

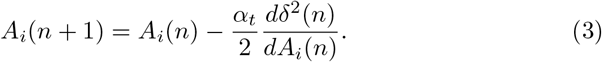

Solving the derivative yields

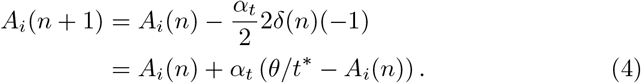

Since the organism only has access to the psychological time given by its internal timing mechanism, and not the physical time t, we assume that an internal estimate for *t* is formed by dividing the current pacemaker count by the current slope, *t* = Ψ_*i*_(*t*)/*A_i_(n)*. Substituting this estimate into equation (4) we get:

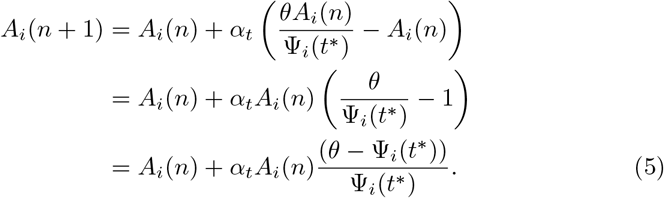

Hence, the update rule for slope *A_i_* to be applied at target time *t*^∗^ (the end of the trial or of the interval being timed) is

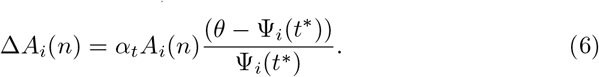

Equation (6) is the slope update rule we use. Note that n above is indexing the number of occurrences of a specific interval that the timer is timing. These intervals may be the duration between CS onset and US onset (the usual ‘trial’ in delay conditioning for example), but they may be any other salient time interval such as CS or intertrial duration. Figure 2 shows timer slope adaptation during three timing scenarios: timing a novel stimulus (row 1), timing a long-short change in stimulus duration (row 3), and timing a short-long change in stimulus duration (row 5).

**Figure 2:**
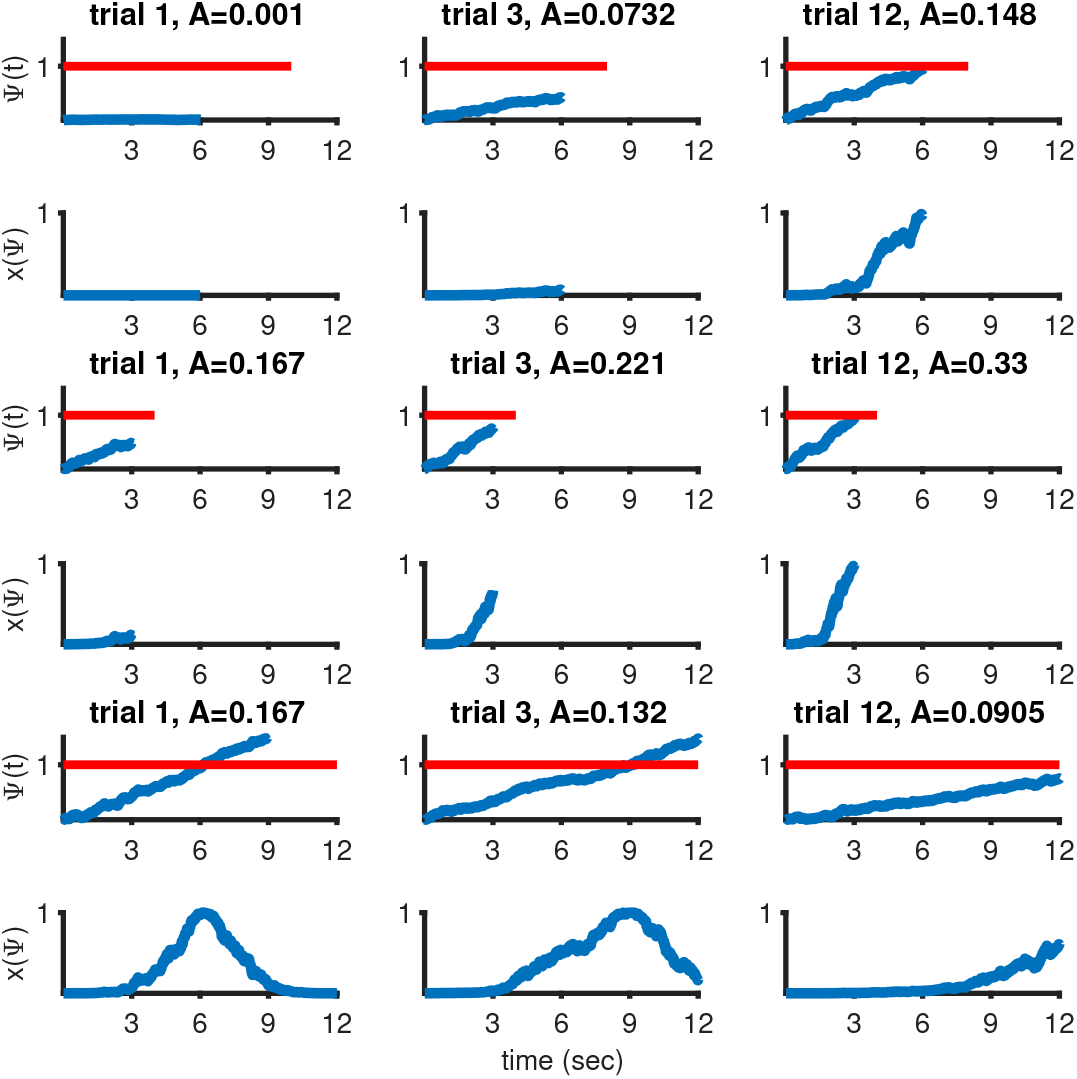
RWDDM timer and CS representation during three 12-trial timing scenarios. Top two rows: timing a novel 6 second stimulus. Timer starts with a low baseline slope (*A* = 0.001) on trial 1 and gradually adapts over training to reach approximately the required slope. Middle two rows: stimulus duration change from 6 to 3 seconds. Bottom two rows: stimulus duration change from 6 to 12 seconds. Parameters: *α* _*t*_ = 0. 215, *θ* = 1, **σ** = 0. 25, m = 0. 15.

In the top row of figure 2 and throughout the paper we assume that the initial value of slope *A* for a novel stimulus is so low as to overestimate the stimulus duration. This overestimation will only last for a few trials, the number of which can be made arbitrarily small by choosing a high adaptation rate *α_t_*. Alternatively, it would be possible to use a very high initial value for *A* so as to underestimate the stimulus duration. However this alternative does not seem neurophysiologically plausible as the brain would need to keep a pool of neurons firing very rapidly as its ‘standby’ timer.

In TDDM, timescale invariance arises from the nature of the noise in the accumulator. After repeated training, say in delay conditioning with a CS of fixed duration, equation (6) will converge to a value of *A_i_* which will make the accumulator reach the threshold value *θ* at the time of stimulus offset, but only on average. In some trials the accumulator will reach the threshold sooner, in which case the organism will underestimate the stimulus duration. In other trials the accumulator will reach the threshold later, causing overestimation. The variability of this time estimate relative to the mean is given by the coefficient of variation (CV). It has been well established experimentally that the CV of time estimates in humans and other animals is approximately constant over a wide timescale (Gibbon, 1977; Gallistel and Gibbon, 2000; Allman et al., 2014). The CV of TDDM’s time estimate is (see equation 3 in Luzardo et al., 2017)

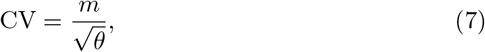

which depends only on the choice of threshold *θ* and noise factor *m*. As these are constant, the CV of TDDM’s time estimate is also constant. Note that because the timer adapts its slope gradually, if the duration of a CS is changed, CV measurements will only match the one given by equation (7) after the slope has finished adapting. The number of trials to adaptation will vary depending on the adaptation rate *α_t_*.

We substitute the presence representation used in the original RW model by a Gaussian radial basis function. Its input is provided by the TDDM accumulator:

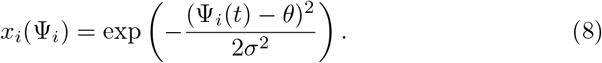

This representation may be interpreted as the receptive field of time-sensitive neurons that read the signal coming from the accumulator neurons. Their receptive fields are tuned to the accumulator threshold value *θ*. The bottom right panel in figure 1 shows the representation for CS_*l*_ generated from the input provided by the timer on the left. Note how *x_l_* reaches its maximum value at the same time that Ψ_*l*_ crosses the threshold at 1. Figure 2 shows x(Ψ) adapting in the three different timing scenarios explained previously. As can be seen, *x_i_* is a dynamic representation of CS_*i*_ that adapts to the temporal information conveyed by the stimulus. Other representation shapes could be used, like a sigmoid for example, but a Gaussian is mathematically simple and has been used before by at least one other timing model (MS-TD, Ludvig et al., 2008).

We follow Gibbon (1977) and Gibbon and Balsam (1981) in assuming that time sets the asymptote of learning, λ, in equation (1). They were led to this hypothesis by investigating CR timing in fixed interval conditioning schedules, a type of delay conditioning. After enough training in this procedure, subjects begin responding some time after CS onset, with a slow rate at first which then increases rapidly until it reaches asymptotic level some time before reinforcement delivery. Gibbon (1977) proposed that subjects make an estimate of time to reinforcement which is used to generate an expectancy of reinforcement. The expectancy for a particular CS_*i*_ with duration *t*^∗^, *h_i_*, was hypothesised to be hi = *H/t^∗^*, where *H* was a motivational parameter which was assumed to depend on the reinforcing properties of the US. The reinforcing value of the US is thus spread evenly over the CS length. It was assumed that this expectancy would be updated as time elapsed during the CS, such that *h_i_ (t) = H/*(*t^∗^-t*). Hence, expectancy would increase hyperbolically until the estimated time to reinforcement *t* = *t*^∗^. Responding would reach asymptotic level when the expectancy crossed a threshold value *h_i_* (*t*) = *b*.

Here we will not use Gibbon’s concept of expectancy update. A similar role is fulfilled by the TDDM accumulator in our formalization. But we hold on to his argument that the reinforcing value of the US is spread over the CS length. Within the Rescorla-Wagner modelling framework, Gibbon’s expectancy value may be interpreted as setting the asymptotic level of learning in equation (1), namely *λ* = *H/t*^∗^. Under this interpretation, *λ* may be said to implement hyperbolic delay discounting of rewards. Similarly to the argument used above in the derivation of the slope update rule, we use the psychological time estimate from TDDM in place of the physical time *t*^∗^, such that *t*^∗^ = Ψ*_i_* (*t*^∗^)/*A_i_ (n)*. The value we use is then 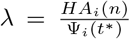. Another possibility would be simply *λ* = *H A_i_(n)*. Both alternatives yield the same asymptotic value, but *H A_i_(n)* converges gradually (with the rate set by *α_t_*) whilst 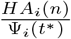 immediately. Our version of equation (1) for updating associative strength then becomes:

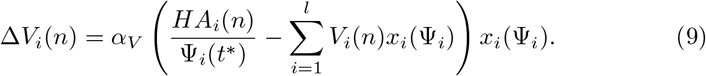

In the trial-based RW model, equation (1) is applied at the end of a ‘trial’, which is usually taken to be the event starting at CS onset and ending at US delivery. We follow the same practice here and apply equation (9) at the end of a trial, i.e. at US delivery. Note that because *x_i_* (Ψ_*i*_) is a dynamic CS representation, its activation (or strength) level at the end of the trial will vary from trial to trial, as can be seen in figure 2. Equation (9) is applied using the activation level of *x_i_* (Ψ*_i_*) current at the end of the trial.

We assume that real-time responses to a CS_*i*_ are emitted according to the product of its associative strength *V_i_* (*n*) and representation *x_i_* (Ψ_*i*_), that is, it is the output of the summing junction in figure 1:

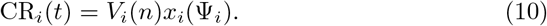

Equations (2), (6), (8), (9), (10) fully define the basic model. Its six free parameters are: *m, *α*t, θ, *σ*, *α*V, H*.

### 2.1 Relationship with Other Models

Among the theories capable of providing an account of both timing and conditioning, arguably four stand out for their scope or influence. They are CSC-TD, MS-TD, LeT and MoT.

TD has been developed primarily as a learning model, without the explicit intention of addressing timing. It may be visualized as a real-time rendition of the RW rule. Its basic learning algorithm, is given by:

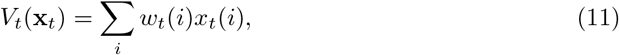

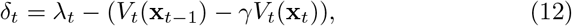

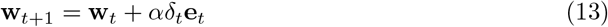

where *V_t_* is the US prediction at time t, formed by a linear combination of the weights *w*(*i*) and the CS representation values *x*(*i*). This update algorithm is performed at each time step, and not only at the end of a trial like RW and RWDDM. Another important difference is that equation (12) computes a difference between the current US value and the temporal difference between predictions. Hence, *δ_t_* > 0 if the US is higher than this temporal difference in prediction, and *δ_t_*<0 if the US is lower. The constant 0 < *γ* < 1 is termed a discount factor. Equation (13) updates the weights for the next time step. The vector et stores eligibility traces, which are functions describing the activation and decay of representations x*_t_*. The three most common eligibility traces used are: accumulating traces, bounded accumulating and replacing traces. These three types accumulate activation in the presence of the CS and discharge slowly in its absence, the first accumulates with no upper bound, the second only until the upper bound and the third is always at the upper bound whilst the CS is present (Sutton and Barto, 1998, pp. 162-192).

The richness of TD’s timing account relies on the choice of CS representation x. The Complete Serial Compound representation (CSC, Moore et al., 1998) postulates one CS element *x*(*i*) per time unit of CS duration. Each element is only switched on at its activation time unit, and then decays afterwards following its choice of eligibility trace *e*(*i*) (usually an exponential decay function). This componential representation, which increases in size linearly with CS duration, should be contrasted with RWDDM’s molar representation (equation (8)) which requires only one element. CSC may be called a time-static representation, whilst RWDDM is a time-adaptive representation, with a rule to change its structure based on a change in time (equations (6) and (8)). CSC-TD also lacks any mechanism to explain timescale invariance of the response curve, which is present in RWDDM. A modification of CSC has recently been developed, the Simultaneous and Serial Configural-Cue Compound (SSCC, Mondrago´n et al., 2014). SSCC-TD formalizes the idea that when multiple stimuli are presented together in time, a configural cue–a novel stimulus that is unique to the current set of present stimuli–is formed. SSCC follows on the CSC representation, but, unlike any other TD model, it allows for the representation of compounds and configurations of stimuli. Because SSCC-TD is a real-time model, it also allows for the simulation of CR timing during compounds and configurations. However, its approach to timing is still the same as CSC, i.e. it breaks down the stimuli into a series of elemental units which are activated in series. Therefore, with respect to timing only we will consider SSCC to belong to the family of CSC representations.

The Microstimuli representation (Ludvig et al., 2008,0) introduced a more realistic description of time. Unlike CSC, it uses a fixed number of elements *x*(*i*) per stimulus. The ith microstimulus is given by:

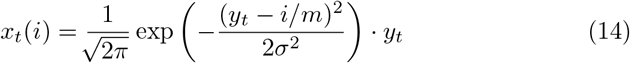

where *m* is the total number of microstimuli, *y* is an exponentially decaying time trace set at 1 at CS onset. It will be noted that a microstimulus is a Gaussian curve modulated by the decaying trace *y_t_*. The set of microstimuli generated by the CS will then give rise to partially overlapping Gaussians, with decreasing heights and increasing widths across time. The fact that only a fixed number of microstimuli are required per CS is an improvement to the potentially large numbers of elements in CSC. The MS representation tries to capture the idea that as time elapses, the stimulus leaves a more diffuse and faint impression. However, even though it is more realistic than CSC, it still lacks a mechanism to produce exact timescale invariance.

Learning to Time is primarily a theory of interval timing which can also account for some aspects of conditioning. Here we will deal with its most recent version in Machado et al. (2009), which differs somewhat from the earlier version in Machado (1997). Its CS representation resembles CSC in postulating a long series of elements (or states) that span the whole stimulus duration. Unlike CSC, it transitions from state to state at a rate that varies from trial to trial, and that is normally distributed. Hence, time during a trial is represented as a noiseless linear increase from states *n* = 1, 2, 3,… (one per time-step) at a fixed rate. This linear time representation resembles the linear accumulator in RWDDM, except that the latter has noise built into the linear accumulator, whilst LeT assumes noise only at the intertrial level. Each state *n* is associated with the US via an associative link. At the end of a trial, the strength *w* of these links are updated as follows:

- For the active state at reinforcement, *n*^∗^, the update rule is 

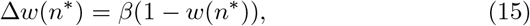
 where *β* is a constant.

- For inactive states, *n* < *n*^∗^, the update rule is

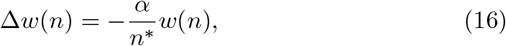

where *α* is a constant.

- For states that did not become active during the trial, *n* > *n*^∗^, the rule is

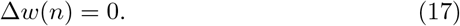

Note that unlike RWDDM’s associative update rule, equations (15) to (17) do not include a summation term. This places a severe limitation on the ability of LeT to deal with compound conditioned stimuli. LeT’s strength lies on its being able to explain timescale invariance of the response curve. Machado et al. (2009) showed that it is possible to derive timescale invariance using only the assumption of intertrial normality of state transition rate. Finally, LeT assumes that responses are emitted at a constant rate if the current active state has associative strength *w*(*n*) greater than a threshold *θ*. The fact that responding depends on the associative strength of the current state, and that this strength only changes with US associations, prevents LeT from accounting for changes in timing that are not related to US occurrence. For example, there is evidence that animals learn the timing of a preexposed CS (Bonardi et al., 2016) and are sensitive to changes in timing during extinction (Guilhardi and Church, 2006), two situations that do not involve the occurrence of a US.

Modular Theory is another primarily timing theory that can also deal with some aspects of conditioning. It treats the onset of a stimulus as signalling a time expectation to reinforcement. Its time representation *T* is, like LeT, an accumulator that increases linearly with time *t, T = ct*, where c is a constant. When reinforcement is delivered the current reading from the accumulator is stored in what is called *pattern memory*. Pattern memory is updated at each trial *n* according to

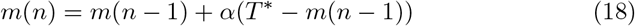

where *α* is a learning rate and *T*^∗^ is reinforcement time. Equation (18) may be contrasted to (6) from RWDDM. The main difference is that pattern memory in MoT stores a moving exponential average of intervals, whilst the slope in RWDDM stores a moving exponential harmonic average of intervals. However, both models are similar in that they can potentially time the occurrence of any event, not only rewards. MoT’s pattern memory and RWDDM’s slope can be made, for example, to adapt to mark the end of stimuli that are not necessarily paired with a reward.

A stochastic threshold *b* is used to mark response initiation. The threshold distribution is set so as to yield timescale invariance of the response curve. Its mean, *B*, is a fixed proportion of the value in pattern memory, *B* = *km*(*n*), where *k* is the proportionality constant, and its standard deviation is γB, where *γ* is the coefficient of variation of *B*. Hence, the coefficient of variation of the threshold, i.e. of response initiation, is constant for all intervals, which is the timescale invariance of the response curve. RWDDM derives timescale invariance of response curve from noise in the accumulator (equation (2), not from the threshold.

This account of time from MoT is an instantiation of Scalar Expectancy Theory, arguably one of the most successful timing models to date. Being a purely timing theory, SET does not address associative learning directly, so it does not have a rule for changes in association between stimuli. MoT bridges this gap by adding a rule to update what is termed *strength memory, w(n)*. Strength memory holds the associative strength between stimulus and reinforcement. The rule consists of a linear operator:

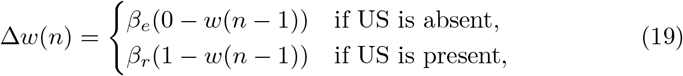

with *β* a constant that can determine different rates of update for acquisition (βr) and extinction (βe). Equation (19) may be compared with (9). Note that, unlike RWDDM, equation (19) does not contain the summation term from RW based rules.

MoT also includes a rule for response rate that is more realistic than RWDDM’s given by (10). It is partly derived from an empirical analysis of real-time responding in animals. We refer the interested reader to Guilhardi et al. (2007) for a fuller description. We will only mention here that MoT generates a two-state response pattern, low and high. The transition between states is determined by the crossing of threshold *B*, and the high state is proportional to strength memory *w (n)*.

Other theories exist which are similar in scope to CSC-TD, MS-TD, LeT and MoT. Two notable examples are the Componential version of the Sometimes Opponent Process model (C-SOP, Brandon et al., 2003) and the Adaptive Resonance Theory-Spectral Timing Model (ART-STM Grossberg and Schmajuk, 1989). C-SOP builds a CS representation based on two sets of elements, or components, one that includes elements activated as a function of time and another whose elements are randomly activated. Associative strength for each element is updated using the standard trial-based RW rule. Simulations in Brandon et al. (2003) have demonstrated that C-SOP can produce some degree of timescale invariance. ART-STM is a neural net with an input layer and one hidden layer, which allows it to explain nonlinear conditioning phenomena (such as negative pattern) that a single-layer RW neural net cannot. It employs a CS representation that is very similar to the microstimuli used in MS-TD, so it also shows a degree of timescale invariance. Other theories could be mentioned (for two influential examples see Buhusi and Schmajuk, 1999; McLaren and Mackintosh, 2000,0) but we will limit the analysis to CSC-TD, MS-TD, LeT and MoT for two reasons: a) these four models collectively embody most of the conditioning and timing mechanisms used in modelling these areas, and b) our goal here is not to provide a comprehensive review, but rather focus on the mechanisms that are shared by our proposed model and the others.

Table 1 summarizes the main mechanisms/features of the models described above. In terms of the type of time representation, it may be observed that the models fall roughly into two categories: (a) those that employ a chain of units or states activated sequentially (CSC-TD, MS-TD, LeT), and (b) those that employ an accumulator (MoT and RWDDM). Those in category (b) may be considered more economical both computationally and biologically, as they don’t require a number of units that increase with time. In terms of what the representations can time, two categories may be discerned: (a) those that time only rewards (CSC-TD, MS-TD and LeT), and (b) those that can time any stimuli (MoT and RWDDM). Models in category (b) have more flexibility to create a temporal map involving all stimuli present, including those not signalling reward. In terms of timescale invariance, the models are basically divided between those that can account for it (MS-TD, LeT, MoT and RWDDM) and the one that cannot (CSC-TD). Finally, in terms of the type of associative learning rule used, models are divided between those that use a RW-type rule (CSC-TD, MS-TD, RWDDM) and those that use the linear operator (LeT and MoT). The ones that use RW are wider in scope, being able to account for cue-competition phenomena, which form the core of classical conditioning.

**Table 1:**
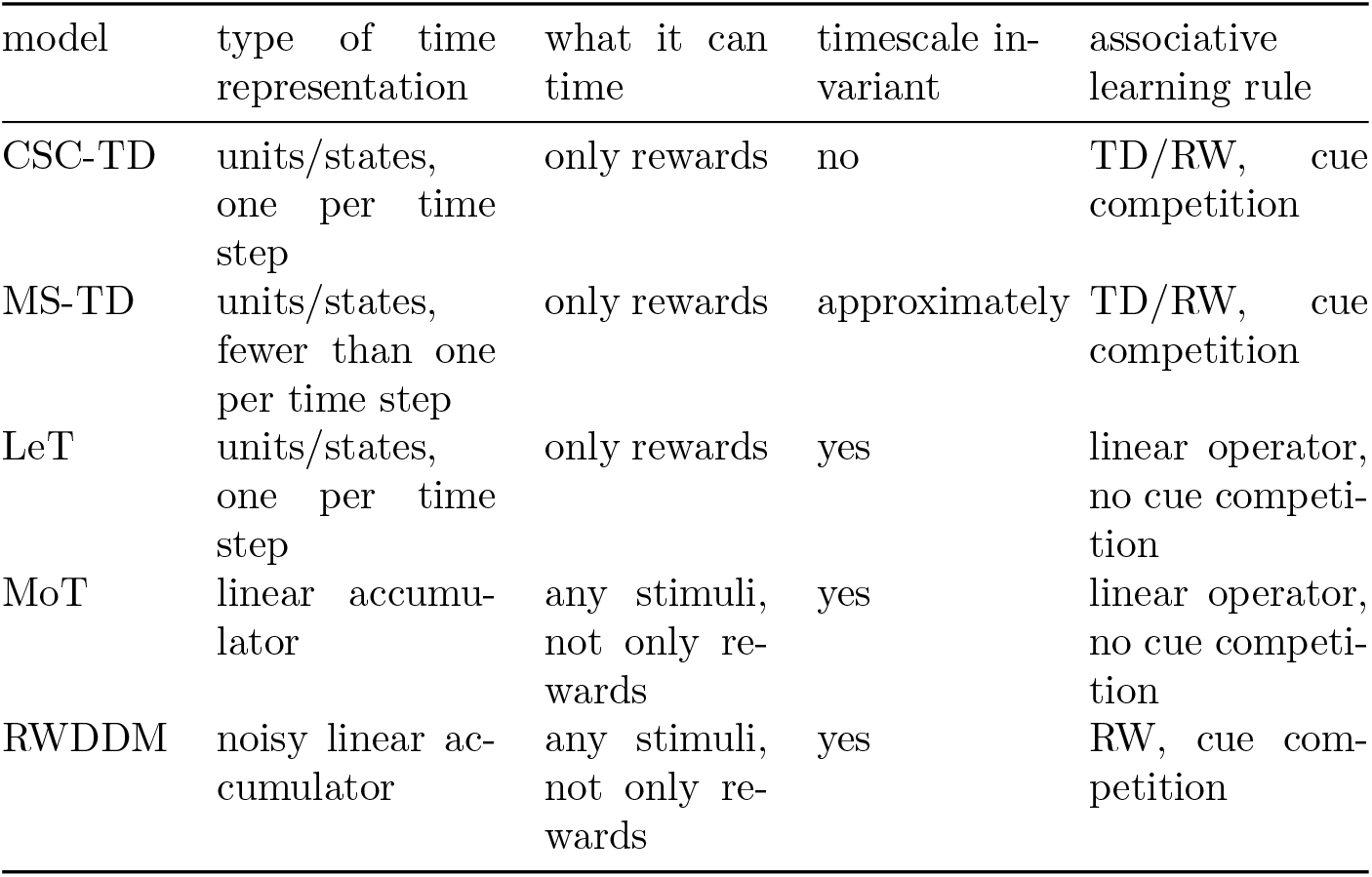
Summary of the main features of the models.

The main innovation of RWDDM over its predecessors is the combination of a noisy linear accumulator for timing with the RW rule for associative learning. As table 1 shows, linear accumulator theories are the only ones in our sample of the models that can fully account for timescale invariance. But because they rely on the linear operator rule, they cannot account for cue-competition and other compound stimuli phenomena in conditioning. Therefore RWDDM extends the application of the linear accumulator to compound stimuli, covering a wider range of conditioning phenomena.

**Table 2:**
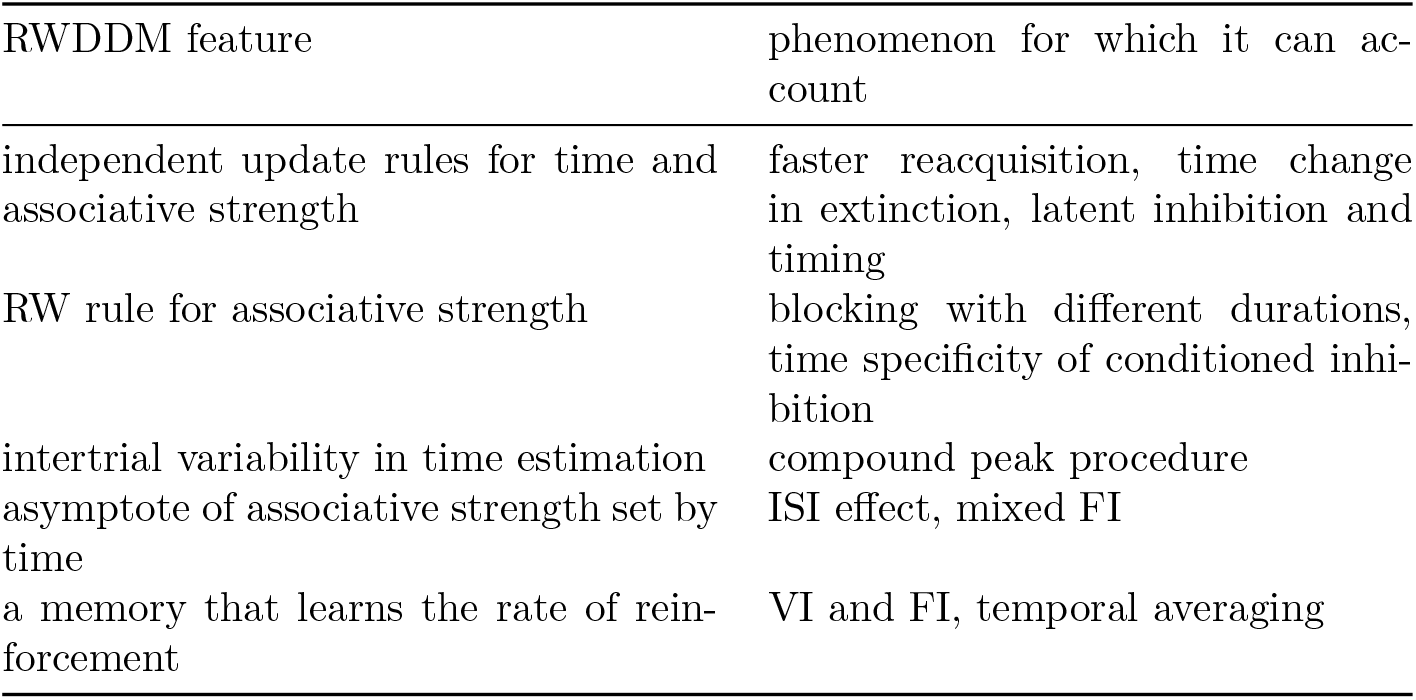
Model features and the experimental findings they can explain.

In summary, the model we propose is, to the best of our knowledge, the only one that unites the flexibility, computational economy and timescale invariance of the linear accumulator as a time representation, to the RW associative learning rule, which accounts for many more conditioning phenomena than the linear operator. In the next section we evaluate the models against a number of phenomena in conditioning and timing.

## 3 Results

The long history of experimental work in classical conditioning has allowed the discovery of a rich variety of phenomena–a recent review (Alonso and Schmajuk, 2012) has catalogued approximately 87. This forces theorists to be selective when deciding which phenomena to simulate when presenting a new model. We searched the literature for phenomena that could test each feature of the model. Table 2 lists the main RWDDM features, together with the corresponding phenomena found in the literature that can test each.

Table 3 contains the design for each simulation performed with the model. The model parameters used in all simulations were kept almost constant but in some cases a few adjustments were found necessary to obtain a better agreement between model and data. We report their values in each simulation below. The time-step was the same for all simulations: *∆t* = 10 msec. Simulations were performed using MATLAB version R2016b. The code to generate the figures in each result section is available as supplementary material.

**Table 3:**
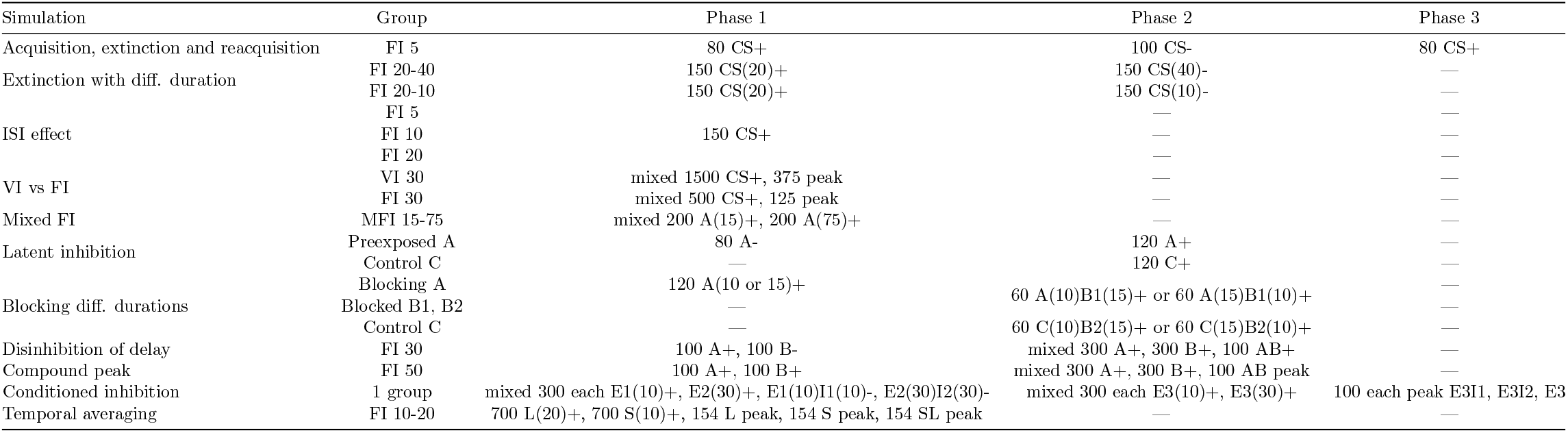
Simulation designs.

### 3.1 Faster reacquisition

A conditioned response emerges gradually over the course of several trials where the CS signals the arrival of a US. If a measure of CR strength (such as rate or magnitude) is plotted against the number of trials, the shape and rate of this acquisition curve will depend largely on the CR and organism, but it usually follows a negatively accelerated curve (Pavlov, 1927; Kehoe and Macrae, 2002). Pavlov (1927) believed timing of the CR would emerge only later in acquisition, through a process he described as *inhibition of delay* whereby the initial part of the CS would become inhibitory. Recent and more detailed analyses suggest that an estimate for the time to reinforcement is acquired very early in training, possibly even after one or two trials, although the expression of such estimation may not be observable until later in training (Holland, 2000; Ohyama and Mauk, 2001; Balsam et al., 2002; Drew et al., 2005).

If the CS no longer signals reinforcement, CR strength gradually decreases over the course of these extinction trials, until it finally disappears. If the CS is made to signal the US again, the CR returns, a process that is called reacquisition. It is a consistent finding that reacquisition is faster than acquisition (Ricker and Bouton, 1996; Guilhardi et al., 2007; Kehoe and Macrae, 2002, p. 185).

Learning is loosely defined as an enduring change in behaviour as a result of experience. Acquisition of a CR is the most basic demonstration that classical conditioning is a form of learning. As such, all classical conditioning models provide an account of it.

#### 3.1.1 Simulations

Figure 3 (top left panel) shows a plot of RWDDM’s associative strength as given by equation (9), in a simulation of acquisition and extinction. Acquisition consisted of 80 presentations of a 5-sec CS followed by reinforcement, after which there were 100 extinction trials where *H* was set to zero. The simulations match with experimental data from acquisition and extinction (bottom left panel of figure 3). The simulated acquisition curve asymptotes around the theoretical value given by setting *∆V* (*n*) = 0 in equation (9) and solving for *V*, yielding

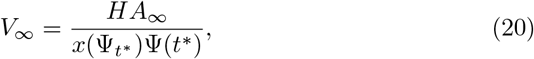

which in this particular case is *V_∞_* ≈ 1, since *H* = 5, *A*_∞_ ≈ 1/5, Ψ_*t∗*_ = Ψ(*t*^∗^) ≈ 1, *x*(Ψ_*t*∗_) ≈ 1, where *t*^∗^ is the time of reinforcement. Because Ψ(*t^∗^*) is a random variable, *x*(Ψ_*t∗*_) and *V_∞_* are also random variables and their values are reported as approximations to their expected values (but not the actual expected values).

**Figure 3:**
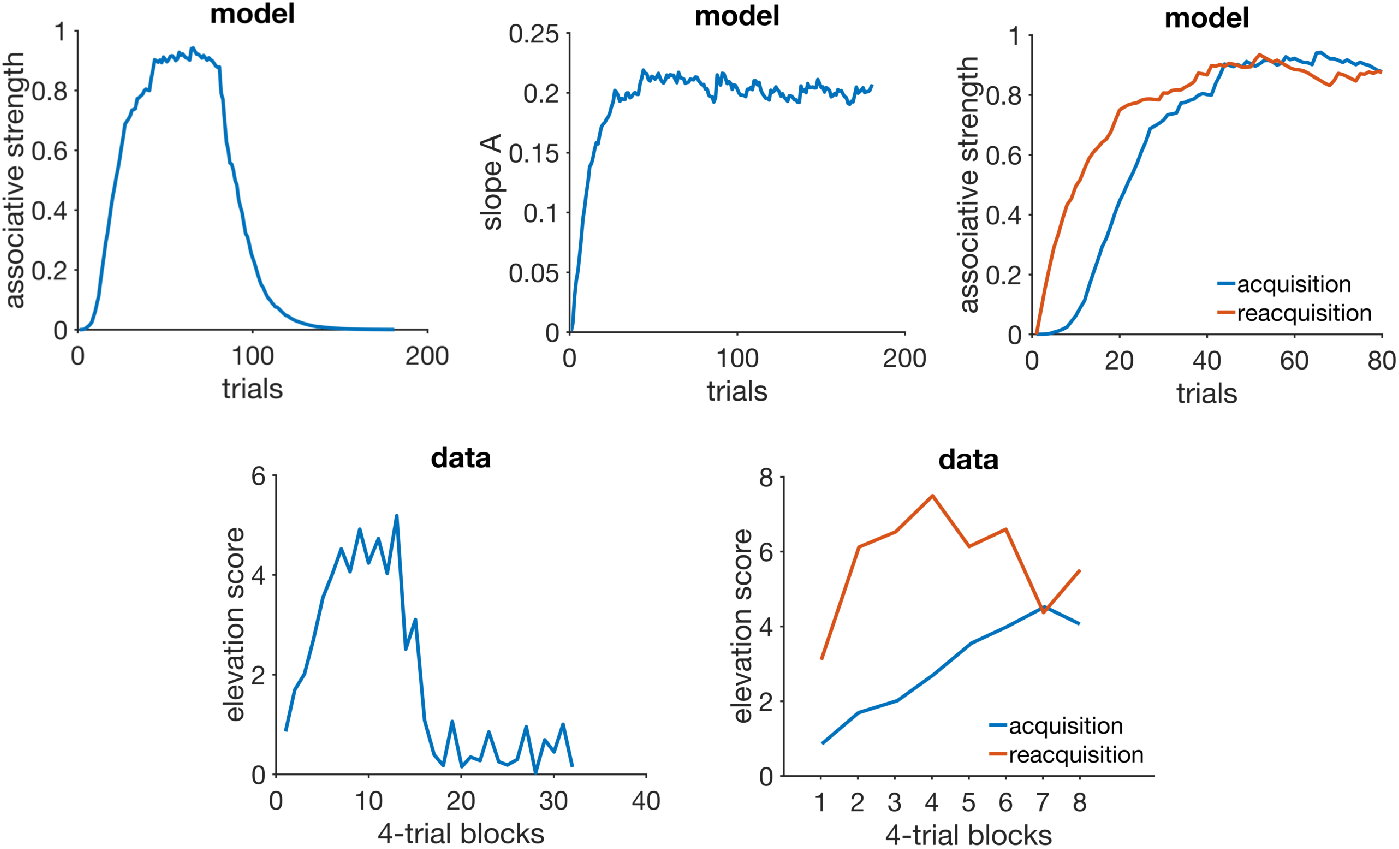
Acquisition and reacquisition. Top left: simulated associative strength *V* in acquisition and extinction. Top middle: adaptation of RWDDM slope A. CR extinction began at trial 80 but has no effect on the RWDDM slope. Top right panel: simulated *V* curves in acquisition and reacquisition. Bottom left panel: response strength data from an experiment in acquisition and extinction, redrawn from figure 1 in Ricker and Bouton (1996). Bottom right panel: data from an experiment in acquisition and reacquisition, redrawn from the top panel of figure 3 in Ricker and Bouton (1996). Model parameters: *m* = 0.15, *θ* = 1, *σ* = 0. 3, *α* _*t*_ = 0. 1, *α*_*V*_ = 0. 1, *H* = 4 in acquisition and *H* = 0 in extinction.

Figure 3 (top middle panel) shows the adaptation of timer slope A given by equation (6). This equation precludes the initial value of A from being zero, so we set it to the very low value of *A*(1) = 10^-6^. We also set the threshold *θ* = 1, which by equation (6) means that *A_i_* (*n*) encodes the exponential moving average of the rate of reinforcement signalled by CS_*i*_. Or, equivalently, 1/*A_i_* (*n*) encodes the moving harmonic average of the intervals since last reinforcement during CS_*i*_. In this simulation, since there is only one US which is delivered always at the same time at CS offset (5000 msec), A converges to A∞ = 1/5000. Note that the value of A does not decline after extinction begins at trial 80. It continues to be updated since the stimulus is still present, even if its presence no longer signals reinforcement.

The top right panel of figure 3 shows the acquisition and reacquisition curves using RWDDM. Reacquisition produced by the model is evidently faster than the simulated acquisition, but not as fast as the reacquisition seen in the data on the bottom left of figure 3.

#### 3.1.2 Discussion

In RWDDM acquisition and extinction of associative strength follow from the same mechanism as RW. The only difference is the noisy stimulus representation *x(_Ψt∗_)*, which induces noise into the acquisition curve. Changes in associative strength and timing are treated independently. In particular, the memory for time encoded by the slope A is not affected by extinction. This leads to a faster reacquisition following extinction. This is because RWDDM’s time-adaptive CS representation *x*(Ψ_*t*∗_) reaches its maximum activation value right from the beginning of reacquisition, since the timer slope A is already tuned to the current CS duration (see equation (8)).

Modular theory (Guilhardi et al., 2007) is another model that treats timing and associative strength separately. It postulates two memories, one for the pattern of reinforcement and another for the strength of the association between CS and US. The pattern memory stores an exponential moving average of the intervals to reinforcement which, like RWDDM, does not change with extinction. However, its strength memory w(n) is updated according to the linear operator rule,

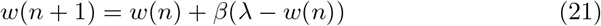

which, unlike RWDDM, does not include a term for a time-adaptive CS representation. Thus, the way MoT accounts for rapid reacquisition is by using different learning rates *β* for acquisition and reacquisition. The same strategy may be employed with the TD and LeT models.

In summary, RWDDM explains reacquisition as the persistence of a memory for time, whilst TD, LeT and MoT explain it as a permanent change in the learning rate for associative strength.

### 3.2 Time change in extinction

When a previously conditioned stimulus is no longer followed by reinforcement, the conditioned response gradually decreases. An important theoretical question for hybrid timing/conditioning models concerns what happens to the timing of responses in extinction. Using the peak procedure Ohyama et al. (1999) found that although the maximum (peak) response rate decreased in extinction, peak time and sensitivity (measured by the coefficient of variation) remained virtually unchanged. Drew et al. (2004) investigated the behaviour on extinction by changing CS duration between acquisition and extinction. Groups where the CS changed to a shorter or longer duration were compared to another where the duration did not change. They found that CS duration had little effect on the rate of extinction, with all groups taking about the same number of trials to achieve CR extinction. However, when the CS used in extinction was considerably longer (4 times) than the one acquired, extinction was facilitated. Guilhardi and Church (2006) performed a similar experiment (experiment 2) and observed that when stimulus duration is changed from acquisition to extinction, the pattern of responding during extinction gradually shifts to the new duration over extinction trials. Following the same procedure, Drew et al. (2017) also used partial reinforcement to slow down the rate of acquisition, and thus observe if response patterns really do shift gradually to the new duration. They confirmed that when CS duration was increased from acquisition to extinction, the within-trial response peak shifted gradually to the right over the course of extinction. When the CS was shortened, the results were not conclusive. Also, when CS duration was changed from training to extinction, the speed of extinction increased, but this appeared to be explained at least in part by the shifting of response patterns.

In summary: a) peak timing and CV are not altered in extinction when using a peak procedure, b) changing the CS duration from training to extinction causes the within-trial response peak to shift to the new duration, and c) changing the CS duration in extinction can speed up extinction, but this may be due to the shifting of the response peak and not to changes in associative strength. These results pose a challenge to the models analysed here. Out of CSC-TD, MS-TD, LeT and MoT, only MoT has a mechanism that would allow it to account for time change in extinction.

#### 3.2.1 Simulations

RWDDM provides an account for these findings as follows. In the case of the peak procedure, the occurrence of the longer peak trials may be considered too infrequent to cause a shift to the longer time. In this case, equation (6) is not applied in peak trials so RWDDM predicts that both slope A and CV will remain unaltered in extinction. In the case of a permanent change in CS duration from acquisition to extinction, the slope update rule is applied and the response peak will shift gradually to the new duration.

We have simulated RWDDM in two extinction conditions, one where the CS presented in extinction was longer than the one acquired (20 sec to 40 sec, short-long) and another where the extinction CS was shorter than the acquired CS (20 sec to 10 sec, long-short). Figure 4 summarizes the main results.

**Figure 4:**
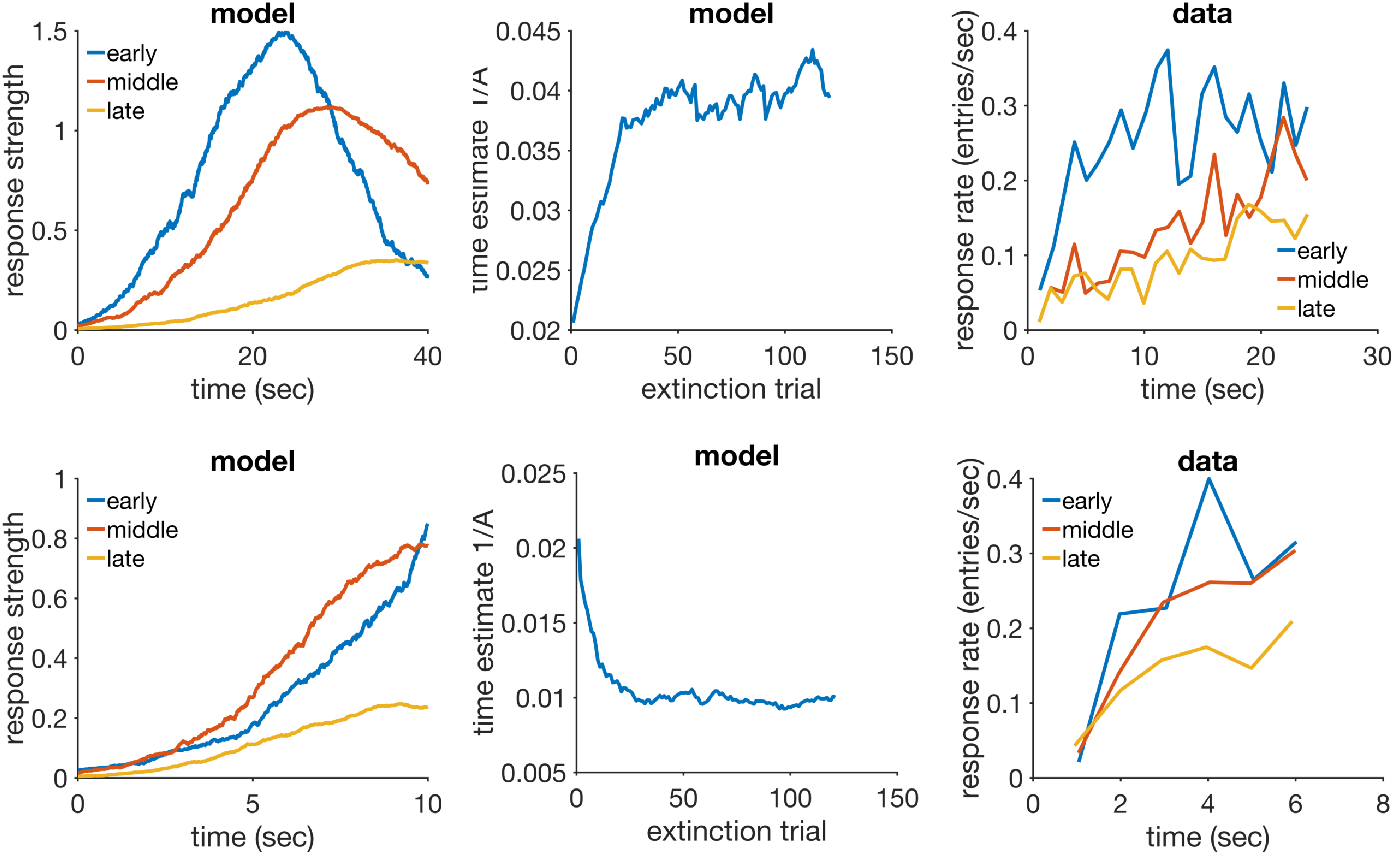
Time change in extinction. Left column: simulated response strength averaged over trials in extinction short-long (top) and long-short (bottom). Middle column: time estimate adaptation of the model during extinction short-long (top) and long-short (bottom). Right column: experimental data from an experiment where the CS duration changed from 12-sec in acquisition to either 24-sec (top) or 6-sec (bottom) in extinction. Data plots redrawn from figure 10 in Drew et al. (2017). Model parameters: *m* = 0.25, *θ* = 1, *σ* = 0.35, *α_t_* = 0.08, *α*_*V*_= 0.09, *H* = 30.

The panels on the left column show response strength during a trial in conditions short-long (top) and long-short (bottom). In the early stages of extinction (early) the response curves peak around the time of US arrival in acquisition (20 sec). This is more evident in the condition short-long (top left) because in the other condition (bottom left) the trial ends 10 seconds before the peak at 20 seconds occurs. Had the stimulus remained on for a full 20 seconds, the response curve in the early stages of long-short would have continued to increase until the 20 second mark. In middle and late extinction the response peak slowly shifts to the new duration in both conditions, and their heights decrease. Compare the simulated curves in the left column of figure 4 to the actual experimental data in the right column. The panels on the middle row of figure 4 show the adaptation of time estimate 1/A in conditions short-long (top) and long-short (bottom). They demonstrate that RWDDM adapts exactly to time change in extinction.

To investigate if the rate of acquisition changes with CS duration, we have plotted the extinction curves for each CS duration in the left panel of figure 5. Decreasing CS duration from acquisition to extinction slightly facilitates extinction, but increasing CS duration markedly delays extinction. However, these are only the *V* values, a theoretical construct that accounts for the associative strength of the stimulus as a whole. Actual behaviour measurements of extinction are based on how much response frequency changes from trial to trial. But response frequency also changes within the trial. As pointed out by Drew et al. (2017), the value obtained for the rate of extinction may be affected by which portion of the CS was measured. To analyse this, Drew et al. (2017) measured response frequency only during the first 6-sec (half the duration of the CS in acquisition) of each CS duration in extinction. We have followed the same procedure and the results can be seen on the middle panel of figure 5. They show a marked delay on extinction when the CS duration was shortened, but not when it was lengthened. Compare these curves with the actual data analysed by Drew et al. (2017) and displayed in the rightmost panel of figure 5. The simulations conflict in part with the same analysis in Drew et al. (2017), which showed no delay on extinction, only facilitation in the case of extending CS duration.

**Figure 5:**
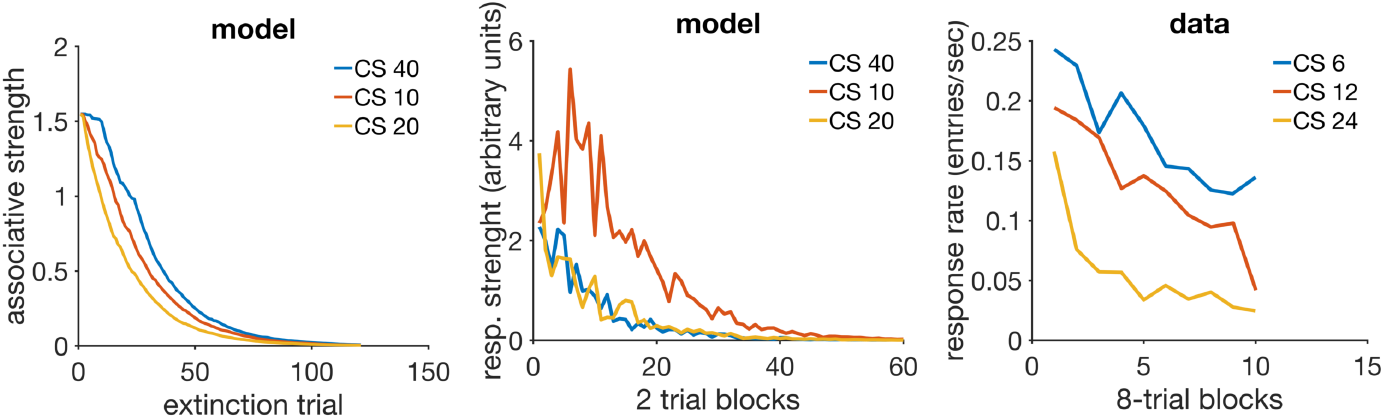
Extinction curves. Left panel: model *V* values for each CS duration in extinction. Middle panel: simulated CR values calculated only for the first 10 seconds of the CS. Each data point is calculated by summing the output of equation (10) over the first 10 sec of each trial, then averaging these trial values two by two, and dividing by 100 to rescale. Right panel: actual CR data for the first 6 sec of the CS in extinction, redrawn from figure 8 (C) in Drew et al. (2017)

#### 3.2.2 Discussion

RWDDM predicts that a change in CS duration from acquisition to extinction will always cause a rescaling of the response curves in extinction. This is largely in agreement with the data. However, RWDDM seems to predict a degree of delay on extinction, whilst the data seems to point to a facilitation of extinction when the CS changes duration. When only the first half of the CS response curves are analysed, the data suggests that extending CS duration in extinction can speed up extinction, whilst RWDDM predicts that shortening CS duration will delay extinction.

RWDDM’s prediction for a delay in extinction following a change in CS duration is due to the shifting of the response curve. At the beginning of extinction, a trial ends either before the CS representation has reached its peak (CS shortening) or after its peak (CS lengthening). This makes equation (9) update with a small value for *x*(Ψ), resulting in a smaller update than with the higher *x*(Ψ) value of the unchanged CS.

As mentioned above, time change in extinction is a difficult phenomenon for the current models to explain. CSC-TD does not have a mechanism to change the peak of responding when a US is not present. Neither does MS-TD or LeT. These models assume that extinction can only weaken existing links between CS and US representations. Because in these models timing usually depends on the sequential activation of these links, changing the CS duration in extinction would not alter the timing but only the magnitude of responding. RWDDM explains time change in extinction because its rule for time adaptation is independent of a change in associative strength. Thus, when the duration changes in extinction, RWDDM’s accumulator slope tracks this change, whilst associative strength decays as a function of US absence. Regarding the extinction facilitation caused by a change in CS duration, none of the models analysed here currently have a mechanism to explain this either.

It would be possible to allow the average rate of state transition in LeT to vary as a function of CS duration, which would cause timing to adapt to the new time in extinction. However, in its latest formulation (Machado et al., 2009) LeT relies on a fixed average rate of state transition to explain timescale invariance. Thus, if the rate is made to change as a function of CS duration, this would break timescale invariance.

As for MS-TD, one interesting modification that would likely allow it to explain time change in extinction is to make the microstimuli themselves time-adaptive. Like RWDDM’s time-adaptive CS representation, the microstimuli could be made to ‘stretch’ or ‘compress’ when stimulus duration shortens or lengthens.

Modular Theory is likely to account for time change in extinction, since its pattern memory for time could be made to update even in extinction. That would shift the response pattern to the new time whilst strength memory, which depends only on US presentation, would decay.

### 3.3 Latent inhibition and timing

When a subject is exposed to repeated and non-reinforced presentations of a stimulus it has never encountered before, this procedure is called preexposure. If reinforcement is subsequently paired with the preexposed CS, the initial rate of CR acquisition is usually lower compared to acquisition to a nonpreexposed stimulus, a phenomenon called latent inhibition (Lubow and Moore, 1959). The asymptotic level of conditioning, however, is not normally affected by preexposure (Lubow, 1989). Latent inhibition is an important representative of a class of phenomena involving latent effects. Collectively, these phenomena demonstrate that something is learned about the stimulus even when it does not signal reinforcement. Therefore, latent inhibition cannot be accounted by the Rescorla-Wagner model, since the theory only applies when there are changes in associative strength.

A question relevant for real-time conditioning models is what happens to timing when a preexposed stimulus is conditioned. To answer this question, Bonardi et al. (2016) used CSs of variable and fixed durations (the variable duration CS had the same mean as the duration of the fixed CS) to vary the temporal conditions between preexposure and conditioning phases. Latent inhibition was observed even when the temporal information from the two phases was different. Crucially, timing, as measured by the response gradient within a trial, appeared to improve in the preexposed CS even when the temporal information was different between the two phases.

As alluded to above, latent inhibition cannot be accounted by the associative learning update rule used in RWDDM, the Rescorla-Wagner. However, we show here that RWDDM is compatible with the Pearce-Hall rule (Pearce and Hall, 1980; Pearce et al., 1982), one of the most widely used models for explaining latent inhibition and other latent learning effects. We demonstrate that this modification maintains the basic framework of the RWDDM, and that it can account for latent inhibition and improved timing with preexposure. None of the other models analysed here can account for latent inhibition without modifications. Improved timing with preexposure could be accounted by Modular Theory, but not by the the current version of the other models.

#### 3.3.1 Simulations

The Pearce-Hall model is basically a rule for adapting the learning rate *α*_*V*_ based on the error δ between the predicted US outcome and the actual US outcome. It was originally formulated by Pearce and Hall (1980) and updated by Pearce et al. (1982). We have maintained equation (9) for associative strength, but changed *α_V_* on every trial *n* according to

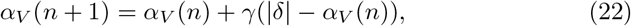

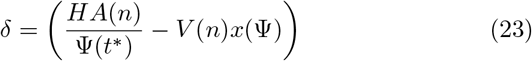

where 0 < *γ* < 1 is a parameter that sets the rate of learning rate adaptation.

Equation (22) is basically the Pearce-Hall rule, except that instead of using 1 as the asymptote of learning we use 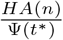.

We simulated latent inhibition with a 5-sec CS. Preexposure consisted of 80 trials of the CS without reinforcement (*H* = 0). The preexposed CS was then reinforced for 250 trials. Figure 6 (top left panel) compares the acquisition curves for the preexposed CS and a control CS in the reinforced trials. The preexposed CS acquisition curve increases at a lower rate than the control CS, the latent inhibition effect (see data from a corresponding experiment at the bottom left panel of figure 6).

**Figure 6:**
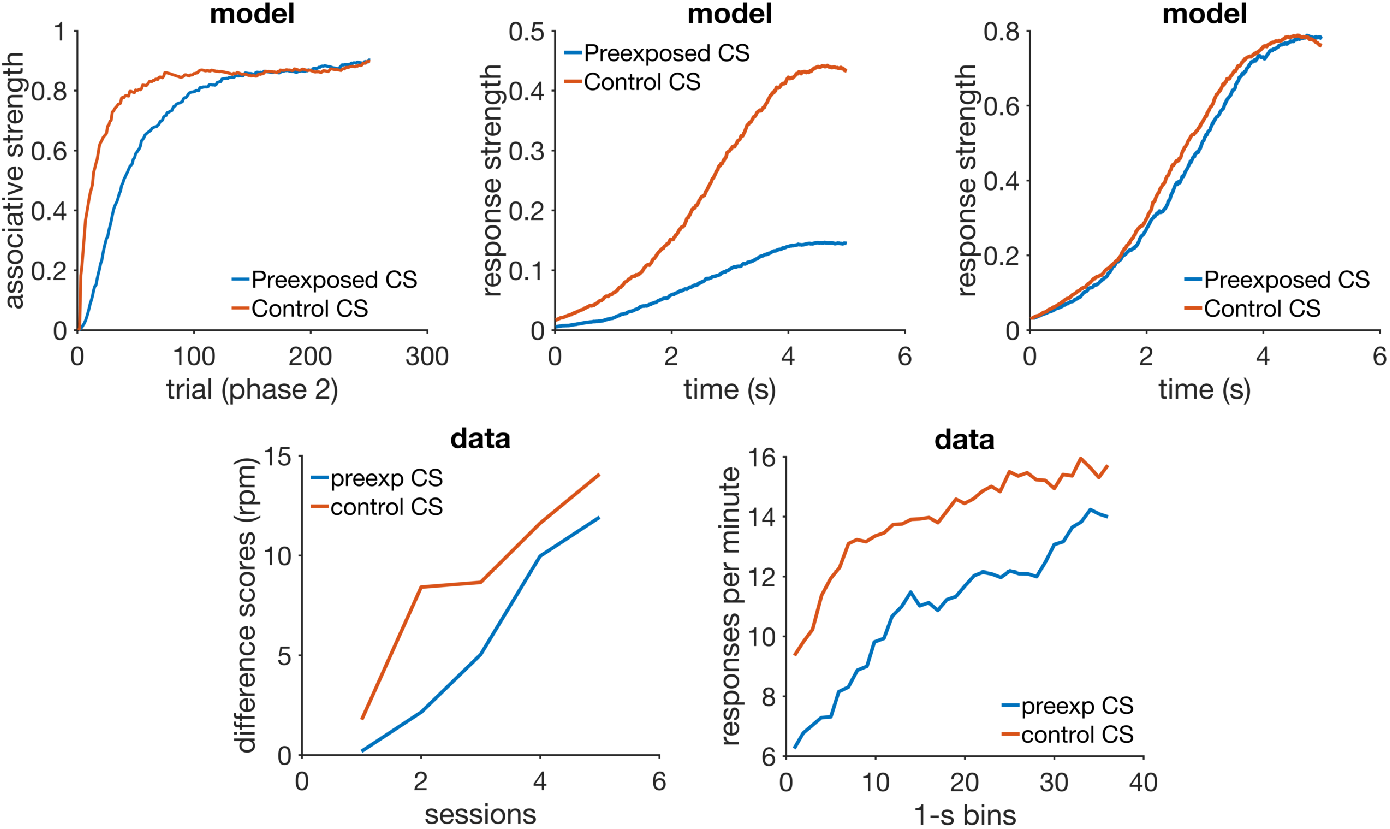
Latent Inhibition. Top row: simulated associative strength in latent inhibition (left), simulated CR averaged over the first 30 trials of conditioning phase (middle), and simulated CR averaged over the last 30 trials of conditioning phase (right). Bottom row: acquisition curves from an actual experiment in latent inhibition (left), and response rate data during the CS (right). Data plots redrawn from figures 1 and 2 respectively in Bonardi et al. (2016). Model parameters: *α_t_* = 0.1, *α_V_* = 0.08, *μ* = 1, *σ* = [0.6-0.35], *m* = 0.2, *H* = 4, *α_PH_* = 0.4, *γ* = 0.03.

Improved timing with preexposure follows directly from the fact that RWDDM adapts its accumulator slope A to the CS duration during preexposure. However, our choice of a Gaussian for stimulus representation does not allow for this change to become visible. Bonardi et al. (2016) demonstrated improved timing by showing that the slope of the response curve from the preexposed CS was higher in the first few trials of acquisition than the one from the control CS (see bottom right panel of figure 6). In general, animal response curves tend to be quite flat during the beginning of acquisition. There is evidence that the response curves appear to change from negatively accelerated to a sigmoidal shape over the course of training (see figure 1 in Meck and Church, 1984, for an example). This means that in the early stages of acquisition, within-trial response frequency increases very early in the trial and then stays at a constant level until the end. As training progresses, the increase in frequency moves slowly to the right, giving rise to the sigmoidal shape that peaks just before the end of the trial. In these cases a higher slope of the response curve would indicate improved timing. But in our model the curves are sigmoidal from start of acquisition, so they will always peak at the end of the trial, even if the timer slope has not adapted to the interval yet, as is the case with a novel stimulus. Therefore, during the acquisition phase of latent inhibition, RWDDM predicts that only the peaks of the response curves will gradually increase over the trials. Because of the learning decrement caused by preexposure, the peak of the control CS will increase faster than the preexposed CS, as the top middle panel of figure 6 demonstrates. The response curve of the control CS will have a higher slope than the preexposed CS, even though the preexposed CS’s timer rate has been adapted to its duration. Hence, the improved timing found in the data is explained by adaptation of RWDDM’s timer slope, but RWDDM’s CS representation cannot make this visible.

We have tried adding an adaptable **σ** in equation (8) so as to decrease the width of the gaussian curve gradually over trials. We chose a simple linear operator rule to adapt the Gaussian width:

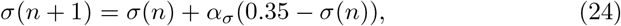

and set *σ*(1) = 0.6 and *α*_*σ*_ = 0.025.

Figure 6 (top middle panel) shows response strength of control and preexposed CSs averaged over the first 30 trials of the conditioning phase. The preexposed CS already shows a clear sigmoidal shape, whilst the control is slightly wider and linear. But the effect is too small to be able to account for the one seen in the data from Bonardi et al. (2016). Towards the end of the conditioning phase the two curves converge (figure 6, top right panel).

#### 3.3.2 Discussion

The simulations show that the model can account for latent inhibition adequately if the Pearce-Hall rule is used (in which case the model would be more appropriately named PHDDM). The PH rule adapts the learning rate *α_V_* based on the level of associative learning between stimulus and reward. When the subject encounters a novel stimulus, it is assumed that *α_V_* has some non-zero starting value 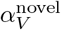, which allows learning in equation (9) to take place. If this novel stimulus does not signal reward, as is the case in the preexposure phase of latent inhibition, **σ** = 0 and equation (22) will simply decay the value of the learning rate across trials until it reaches zero. If at this point the stimulus begins to be followed by reward, **σ** > 0 and equation (22) will begin to raise the value of the learning rate, which in turn will allow equation (9) to begin increasing the value of *V*. Since the increase in the value of the learning rate is gradual, determined by the rate γ, there will be a number of trials in the beginning of the conditioning phase where 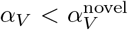, which leads to the initial impairment in the learning curve when compared to the learning curve of a non-preexposed CS, as seen in the top left panel of figure 6.

The separate rule for time adaptation allows the model to account for improved timing after preexposure, but the model cannot make this effect visible even if we allow for Gaussian width adaptation. In view of this it seems more likely that a two-state CS representation may be a better solution. As mentioned above, figure 1 in Meck and Church (1984) suggests that during the initial stages of training a CS representation may be modelled by the following leaky integrator

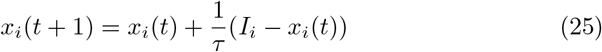

where *I_i_* is the indicator function marking the presence of CS_*i*_, and *τ* a time constant. In the latter stages of training, when timing is expressed, the organism switches to the Gaussian representation given by equation (8). When the switch between representations is made and how abruptly remains to be investigated.

Latent inhibition cannot be accounted by any of the other models analysed here without modifications. Also, models that rely on the US for time adaptation, like CSC-TD, MS-TD and LeT, cannot account for improved timing by preexposure. Modular Theory is the only one that can time any stimulus like RWDDM, so it could account for the improved timing. But it would also need a modification like (22) to adapt its learning rate to account for latent inhibition.

### 3.4 Blocking with different durations

Arguably, the most important compound conditioning phenomenon is blocking. It is part of a class of cue competition and compound phenomena discovered in the late 1960s which challenged the view that conditioning was driven by the pairing, or contiguity, of CS-US. These results suggested that conditioning with compound stimuli was influenced by the reinforcement histories of the elements forming the compound (Rescorla, 1988; Gallistel and Gibbon, 2001). This led to the development of a new generation of models that could account for those findings (Rescorla and Wagner, 1972; Mackintosh, 1975; Pearce and Hall, 1980). The rule we use, the Rescorla-Wagner, provides an explanation for blocking that is based on the summation term in equation (1).

In a blocking procedure a CS is first paired with a US in phase 1 of training. During phase 2 a novel CS is presented in compound with phase 1 CS and paired with the US for just a few trials. Subsequently, when tested alone the novel CS elicits less responding than if it had been trained in compound with another novel stimulus (Kamin, 1968). The previously reinforced CS is said to block the novel CS. The temporal information encoded by each CS has an effect on the amount of blocking observed. Schreurs and Westbrook (1982) varied the ISI in the pre-training and compound phases, and observed less blocking when the durations were different in both phases than when they were the same. Barnet et al. (1993) performed a similar experiment but with forward and simultaneous conditioning varying between phases, and also found that blocking was stronger when blocked and blocking CSs had the same temporal history. Jennings and Kirkpatrick (2006) used compounds where the elements had different durations. They observed that a long blocking CS could block a co-terminating short Cs, but a short blocking CS failed to block a co-terminating long CS (see rows 1 and 3 in figure 7). Amundson and Miller (2008) performed four blocking experiments using trace conditioning. In two of them the blocking CS trace duration changed between phases, and blocking was not observed. In the other two experiments the trace duration was held fixed between phases, and the blocking and blocked CSs were presented serially and not in a compound (see rows 2 and 4 of figure 7). Blocking was observed when the blocking CS followed the blocked CS, but not in the reverse condition.

**Figure 7:**
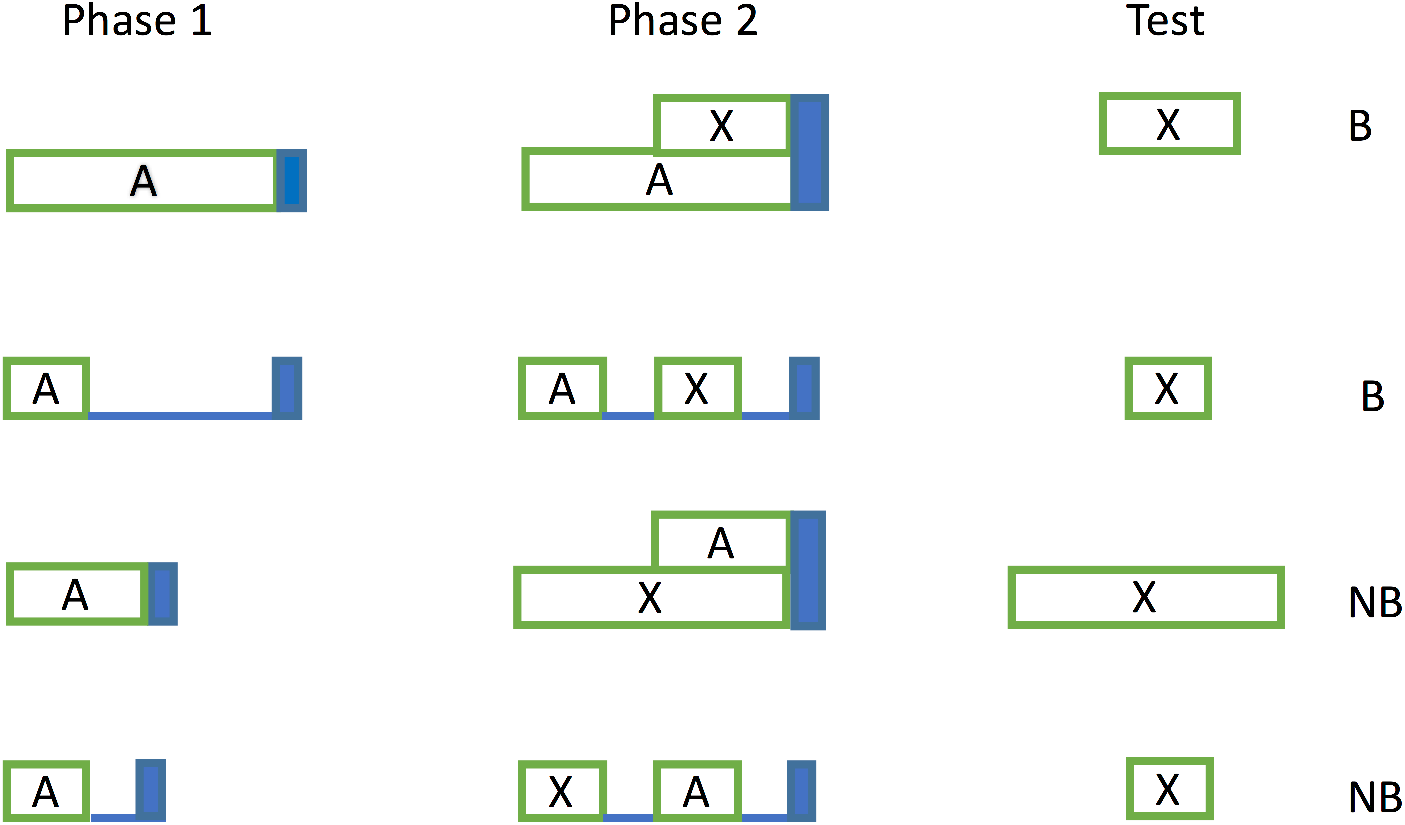
Experimental designs from two blocking experiments. CS X was blocked (B) in rows 1 and 2, and not blocked (NB) in rows 3 and 4. Blue bar indicates US presence.

The studies reviewed above appear to show that changing the ISI of the blocking CS between phases may attenuate blocking. Another finding is the apparent asymmetry of blocking when the ISI of the blocking CS is kept constant between phases. Rows 1 and 2 of figure 7 suggest that a long blocking ISI can block a short blocked ISI. Rows 3 and 4 suggest that a short blocking ISI does not block a long blocked ISI.

As mentioned above, RWDDM can account for blocking because it uses the RW rule. The summation term in equation (1) formalizes the widely held view that a given US can only confer a limited amount of associative strength which CSs must compete for. Different theories exist that take other approaches to blocking (see for example Mackintosh, 1975; Harris, 2006; Stout and Miller, 2007) but among the ones analysed here (for their ability to handle timing also) only CSC-TD and MS-TD are equipped to deal with it. We show next that RWDDM can account for the blocking of a short CS by a long CS, and that by making the reasonable assumption of second-order conditioning it can also account for the lack of blocking of a long CS by a short CS. CSC-TD and MS-TD are also capable of providing an account of both blocking conditions.

#### 3.4.1. Simulations

Because RWDDM is based on the RW rule, it produces virtually the same results as the latter when the CSs have the same duration. Our interest here is to test whether it can reproduce the finding that a long CS can block a shorter CS but a shorter CS does not block a longer one. We performed a simulation following the design in rows 1 and 3 of figure 7. In the first phase a CSA (blocking CS) of duration either 10 or 15 seconds was followed by reinforcement until its associative strength *V* reached asymptote. In phase 2 CSA was joined with a CSX (blocked CS), of either 15 or 10 seconds, in a coterminating compound and followed by US. The top left panel of figure 8 shows the acquisition of associative strength for CSX and its control during phase 2 for the condition CSA-15sec and CSX-10sec. A considerable amount of blocking is observed, matching with the data (bottom left panel).

The top right panel of figure 8 shows the results for condition CSA-10sec and CSX-15sec. In this condition the model diverges considerably from the data (bottom right panel) and predicts that CSX should actually become inhibitory.

**Figure 8:**
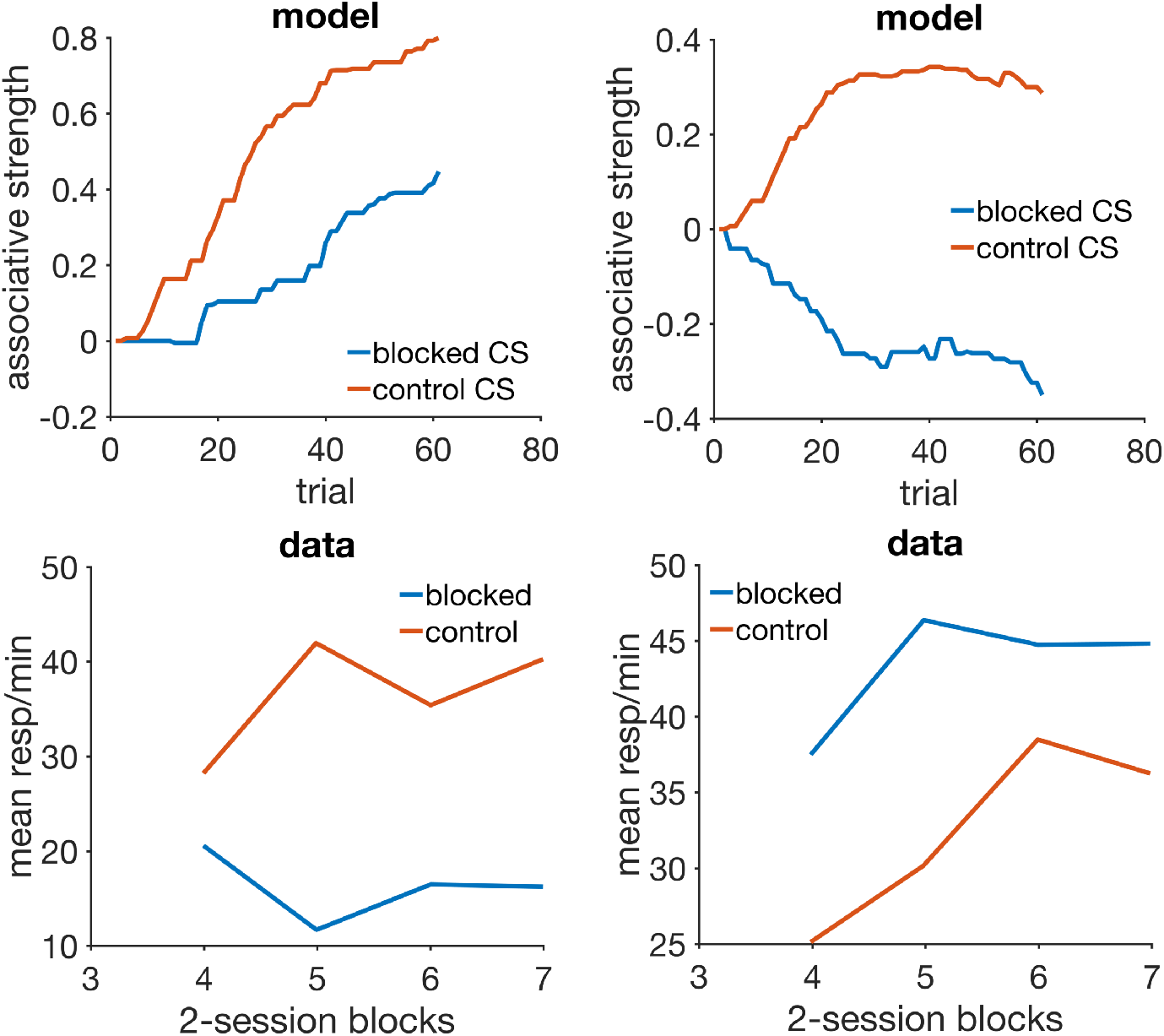
Blocking with different durations. Left column: simulation (top) with a 15 sec blocking CS and 10 sec blocked CS, and animal data (bottom) from an experiment with the same design. Right column: simulation (top) with a 10 sec blocking CS and 15 sec blocked CS, and animal data (bottom) from an experiment with the same design. Data panels redrawn from the top right panel in figure 5 in Jennings and Kirkpatrick (2006). Model parameters: *α_t_* = 0.2, *α_V_* = 0.1, *μ* = 1, **σ** = 0.35, *m* = 0.2, *H* = 10.

#### 3.4.2 Discussion

The blocking and inhibition seen in figure 8 is a result of a discrepancy in the asymptote of learning between the CSs. After phase 1, CSA has associative strength *V_A_ ≈ *H* A_A_*. During phase 2, CSX’s associative strength changes according to:

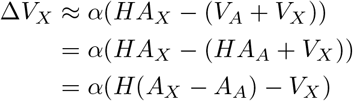

and since (*A_X_-A_A_*) < 0, *V_X_* becomes negative.

However, it could be argued that the short CSA becomes a secondary reinforcer which is signalled by the onset of the long CSX. In this case, the onset of CSX would serve as the time marker for the onset of CSA, and not for the onset of US. Hence, during the first 5 seconds of CSX responding would be under the control of this 5-sec stimulus representation which would not overlap, thus not compete, with CSA’s later representation. It would follow from this account that no blocking would be observed, and that responding during test phase with CSX would peak at the 5-sec mark. This is a testable prediction that, if shown to be the case, could validate RWDDM’s account.

Also note that the time-dependent associative strength asymptote assumed by RWDDM implies that learning during a compound where the elements are of different durations is not stable. In particular, if CSA and CSX are the two elements of the compound phase of blocking, their associative strengths are updated by RWDDM as

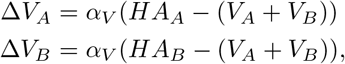

which in the steady state form an inconsistent system of linear equations,

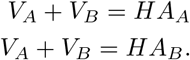

Since the compound phase of blocking only lasts for a few trials, RWDDM could produce the blocking seen on the left panel of figure 8. But if training with the compound was carried out for longer, the *V* values would grow without bound. However, there is evidence that in compounds formed by elements with asynchronous onsets, like in the compound phase of the blocking experiments here, the shorter stimulus comes to control CR timing and there is no summation of associative strengths (Fairhurst et al., 2003). Hence, it appears that with compounded asynchronous CSs, the shorter CS, more proximal relative to the US, comes to dominate and a summation rule like RW would not be applicable beyond the first few trials of training.

A model that is well placed to explain these results is CSC-TD. A long blocking CS will completely overlap a short blocked CS, blocking all units in the blocked CS. But in the case of a short blocking CS, there will be free units in the beginning of the blocked CS which will acquire associative strength, attenuating blocking. Given its similarity, MS-TD would likely produce comparable results. MoT and Let would not be able to account for any type of blocking given their current choice of rule for associative strength. Unlike RWDDM and the TD models, they both rely on the linear operator rule, which antedates the transition to the rules that sum associative strengths in the compounds as mentioned previously. MoT and LeT would need, at the very least, to replace the linear operator by the RW or other equivalent rule to be able to account for blocking and other compound phenomena.

### 3.5 Time specificity of conditioned inhibition

Learning occurs not only when a CS signals the occurrence of a US, but also when a CS signals the omission of a US. It is commonly assumed that the excitation caused by the former is counteracted by an inhibition produced by the latter. This is again formalized by the summation term in the RW rule. Conditioned inhibition is thus one of the phenomena that, together with blocking and other compound phenomena, challenged the contiguity interpretation of classical conditioning.

A conditioned inhibition procedure involves reinforced trials with a CS, say A+, intermixed with non-reinforced trials with a compound AB-. Conditioned responding develops during A+ trials but not during AB-. Hence, conditioned inhibition is a key conditioning phenomenon since it is also a form of discrimination learning.

Conditioned inhibition poses higher technical challenges for a model of learning and timing as responses cannot be directly observed. To assess conditioned inhibition two types of measures are used (Denniston and Miller, 2007): summation and retardation tests. There are different procedures that can generate inhibition, so we refer here specifically to the inhibition produced by alternating A+ with AB-trials. CSA is called a training excitor, and CSB an inhibitor. In summation tests, this inhibitor is then presented together with a different excitor, and the inhibitor is said to pass the test if there is a decrement in responding compared to the excitor alone. In retardation tests, the inhibitor by itself is now paired with the US, and it is said to pass the test if acquisition is slower than with a neutral stimulus. Denniston and Miller (2007) reviewed a series of studies that varied the durations of the training excitor and that between the inhibitor and the training excitor. The studies showed that conditioned inhibition is observed when the temporal relations between training and testing are preserved, and not otherwise.

However, the studies reviewed by Denniston and Miller (2007) used as measure of conditioned inhibition the time to resume drinking (licking suppression) when presented with the inhibitor. Williams et al. (2008) investigated inhibition caused by reinforcement omission in excitatory conditioning, a more direct measure than licking suppression. In their experiments the inhibitor stimulus signalled the omission of one of two USs (at 10 or 30 seconds) that had been associated with the excitor stimulus. Using summation tests they found that the inhibitor would suppress responding only at the specific time of predicted US omission. Retardation tests confirmed that the time of US omission is encoded by the inhibitor.

We show here that RWDDM can account for inhibition and its time specificity. CSC-TD and MS-TD are also equipped to deal with these results. MoT and LeT do not currently have the necessary mechanisms to explain inhibition.

#### 3.5.1 Simulations

We demonstrate time specificity of inhibition with simulations of Williams et al. (2008) experiment. Excitors E1 and E2 signalled reinforcement after 10 and 30 seconds respectively, and inhibitors I1 and I2 signalled US omission after 10 and 30 seconds respectively. During phase 1, E1 and E2 were always reinforced, whilst the compounds E1I1 and E2I2 were never reinforced (see table 3). In phase 2 a transfer excitor E3 was trained on a mixed FI schedule, where in half the trials E3 lasted 10 seconds and in the other half 30 seconds. Phase 3 consisted of nonreinforced peak trials that lasted 90 seconds, a third with E3 compounded with I1, a third with E3I2, and a third with E3 alone. Figure 9 summarizes the results. Responding during E3 alone shows the two peaks characteristic of mixed FIs. As figure 9 shows, the compound excitor and inhibitor inhibits responding only at the time encoded by the inhibitor.

**Figure 9:**
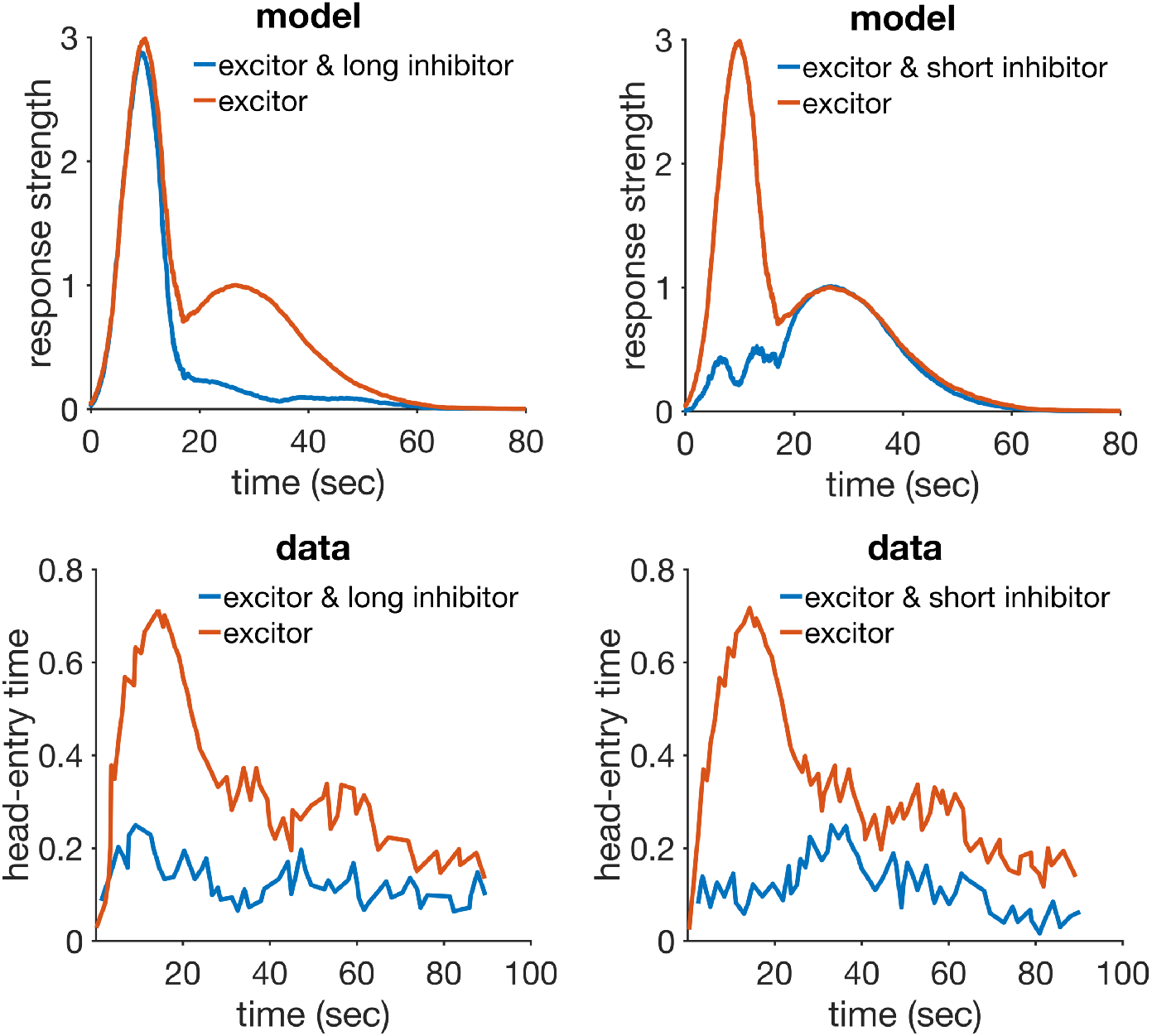
Conditioned inhibition. Left column: simulation (top) and data (bottom) from conditioned inhibition with a long inhibitor. Right column: simulation (top) and data (bottom) from conditioned inhibition with a short inhibitor. Data plots redrawn from figure 4 Williams et al. (2008). Model parameters: *α_t_* = 0.09, *α_V_* = 0.06, *μ* = 1, **σ** = 0.35, *m* = 0.16, *H* = 30.

#### 3.5.2 Discussion

The account provided of inhibition by RWDDM relies on the traditional summation term inherited from the RW rule. Time specificity comes from the inhibitor CS timer being treated just like any other CS timer, except that instead of timing the arrival of the US it times the arrival of US omission.

RWDDM predicts that the representation of an inhibitor CS has the same shape as of an excitor CS. This implies that inhibition is the exact opposite of excitation. This is a testable prediction which the empirical results above provide some validation.

The TD models provide a similar account of these data. Both CSC and MS TD have CS representations that allow for time specificity of US omission. Because the TD relies on the RW summation term, they can account for inhibition. LeT and MoT can also represent such time specificity, but because they rely on the older linear operator rule, they do not have a mechanism to account for inhibition.

### 3.6 Disinhibition of delay and compound peak procedure

The two related phenomena described here are important in that they appear to challenge the summation effect. A common observation is that a compound of two previously conditioned CSs usually produces more responding than its individual components (Rescorla, 1997; Kehoe and Macrae, 2002, p. 204). However, failure to obtain summation is also common (Rescorla and Coldwell, 1995; Pearce et al., 2002), and the precise conditions when it is observed or not is still a current topic of debate (see Harris and Livesey, 2010, for a discussion). Here we consider two cases in which summation was not observed and that RWDDM can offer a possible explanation.

Aydin and Pearce (1995) used an autoshaping procedure to condition pigeons to stimuli of 30 second duration. They observed little or no summation in compound trials, but a response curve with a consistent shift to the left. This earlier start of responding was observed even when one of the components was a neutral preexposed CS. The shift of the response curve to the left was termed disinhibition of delay.

Meck and Church (1984) performed an analogue experiment using the peak procedure. They trained rats to associate a light and a sound (both of 50 second duration) individually to a reinforcement, and then used a peak procedure to investigate what happens to timing in their compound. Like Aydin and Pearce (1995) they also found no summation and a shift to the left in the compound. Furthermore, rats also stopped responding earlier in the compound peak trials.

Taken together, these results appear to show that in some cases summation is not observed, and responding in the compound starts earlier than in the component CSs. One possible explanation for this effect is that the subject fails to recognize the two individual components of the compound, what is known as generalisation decrement. If this is the case then it would be a performance effect, and not a learning phenomenon. We cannot rule this out, but we show that RWDDM’s trial variability in time estimation provides a plausible mechanism to explain this effect. The only other models in our analysis set that can account for this are MoT and LeT.

#### 3.6.1 Simulations

RWDDM is capable of accounting for the earlier responding in compounds by noise in the timer. When a compound formed by CSA and CSB is presented, its two timers Ψ*_A_*(*t*) and Ψ*_B_* (*t*) will run in parallel. However, their rates *A_A_* and *A_B_* will have slightly different values due to noise. This implies that on every compound trial, one timer will be running slightly faster than the other. In contrast, on trials where only one CS is present, the timer will run faster in some trials and slower in others. Therefore, if on compound trials responding is guided by the faster timer, the average response curve for compounds will be shifted to the left when compared to the averaged response curve for a single CS.

Figure 10 shows simulations of disinhibition of delay and compound peak procedure. The figures were constructed by averaging the responses produced by equation (10) over 50 trials. The simulations reproduce in part the anticipation in responding during the compound that is observed in the data in both experiments (see top right and bottom left panels of figure 10). Meck and Church (1984) reported a median peak time of 40±4 seconds for the response curves in compound trials, and 50±3.5 seconds in the individual trials. We ran 15 simulations as the one shown at the bottom row of figure 10, and analysed the peak times produced by each. We found an average peak time of 42±3 seconds in the compound trials, and 47±4 in the individual trials. Both results are within the error bounds in Meck and Church (1984). Aydin and Pearce (1995) did not analyse peak times or shift in the response curves, so we cannot make a quantitative comparison with our simulations.

**Figure 10:**
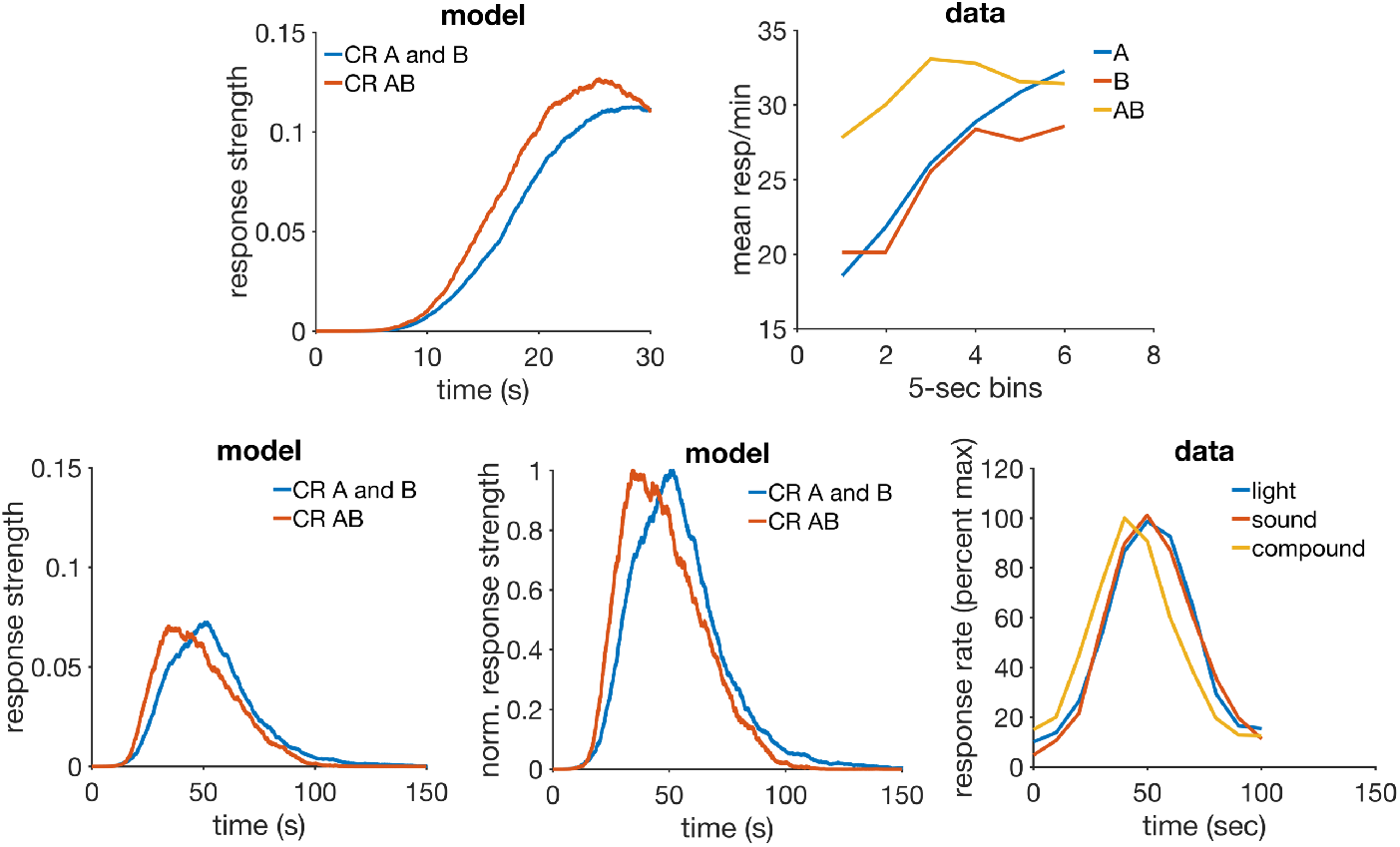
Disinhibition of delay and compound peak procedure. Top row: simulation (left) and data (right) of disinhibition of delay. Bottom row: simulation (left and middle) and data (right) of a compound peak procedure. The middle panel is a normalized (proportion of maximum response strength) version of the left panel. Data plot redrawn from figure 13 in Meck and Church (1984). Model parameters: *m* = 0.25, *θ* = 1, **σ** = 0.18, *α_t_* = 0.75, *α_V_* = 0.1, *H* = 5.

#### 3.6.2 Discussion

RWDDM can offer a good account for the lack of summation and earlier responding in compound trials in the two cases analysed here. It does so by having trial to trial variability in time estimation. However, the model shows a slightly higher maximum response frequency in compounds than in their components (top and bottom left of figure 10) something not observed in the data. This is not the product of summation, but of the slightly different asymptotes of learning in the faster and slower timers in the reinforced trial immediately preceding the peak trial. Our assumption was that in compound trials the timer running faster, with a higher slope A, would be the one guiding responding. When timing adaptation has reached asymptotic levels, the updates on slope A are due to noise in the value of the timer at reinforcement time, Ψ(*t*^∗^). The two slopes, *A_A_* and *A_B_*, will have very similar values. In the reinforced trial preceding the compound peak trial, whichever timer produces a value of Ψ(*t*^∗^) lower than the threshold will have its slope *A* adjusted up by the the slope update rule, likely causing it to overtake the other slope. This slightly higher slope will then be chosen in the peak trial that follows. But the corresponding *V* associated with that timer will have been updated on the previous reinforced trial based on the lower Ψ(*t*^∗^) < *θ* value. Because that is the denominator in *HA*/Ψ(*t*^∗^), the *V* value of the chosen timer will be consistently slightly higher on the compound peak trials.

Other theories that might account for the data in this phenomenon are LeT and MoT. Both theories postulate intertrial variability in timer rate, the same mechanism used by RWDDM to explain this data. TD in any of its current versions lacks a mechanism to explain these data.

### 3.7 ISI effect

The interval between CS onset and US onset is called Inter Stimulus Interval (ISI). In general, measures of CR strength such as response frequency and amplitude decrease with longer ISIs (Smith, 1968; Gormezano et al., 1983; Kehoe and Macrae, 2002). Response timing is commonly analysed by using fixed interval (FI) schedules of reinforcement, which rely on a fixed ISI. It is a well established result that the peak in the response curve decreases with longer FIs (Catania and Reynolds, 1968; Gibbon et al., 1997). However, the entire response curve approximately scales with FI. This is obtained by plotting different FI response curves as the proportion of maximum response strength versus the proportion to FI, a normalization procedure. The resultant normalized curves roughly superimpose (Rakitin et al., 1998; Matell and Meck, 2000,0; Allman et al., 2014). This is sometimes called scalar timing, and it is one of the manifestations of the more general property of timescale invariance.

CSC-TD does not have a mechanism to explain either timescale invariance or the ISI effect. Its more recent development, MS-TD, can approximately reproduce both timescale invariance and the ISI effect. LeT is also a timescale invariant model, but does not appear to show the decrease in response peak as a function of FI. MoT, at least in its earlier version (Kirkpatrick, 2002), can reproduce both the ISI effect and timescale invariance.

#### 3.7.1 Simulations

To demonstrate how RWDDM can reproduce the ISI effect we have simulated a delay conditioning procedure using three fixed interval stimuli. Figure 11 shows RWDDM simulations with FIs 5, 10 and 20 seconds. The top left panel shows within-trial response rate (given by equation (10)) averaged over 50 trials for each FI. The response curves show the same pattern as the data (bottom panel) from the ISI effect: a sigmoidal shape with a maximum that decreases as a function of FI duration. Note that because the curves are averages of 50 trials, the noise is averaged out.

**Figure 11:**
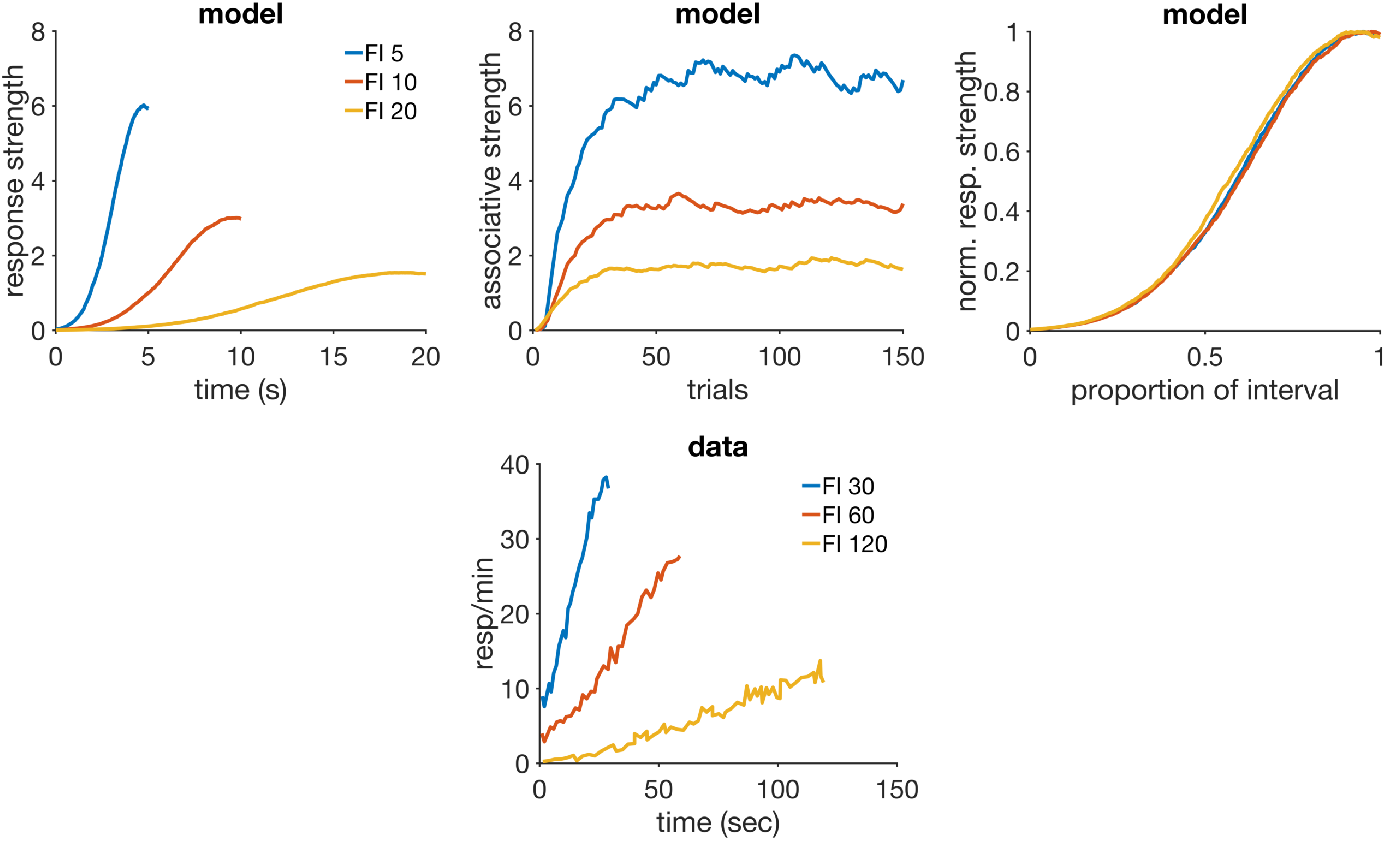
ISI effect. Top row: simulated average response rate during CSs (left), associative strength over trials (middle), and superimposition of response curves (right). Bottom row: average response rate data from an FI experiment, redrawn from bottom right panel of figure 4 in Kirkpatrick and Church (2000). Model parameters: *m* = 0.15, *θ* = 1, **σ** = 0.3, *α_t_* = 0.2, *α_V_* = 0.1, *H* = 5.

The top middle panel of figure 11 shows the associative strength acquisition curves for each FI. Their asymptotic levels are given by equation (20). *V*_∞_ is approximately a linear function of *A_∞_*, the TDDM slope. The different asymptotic levels of associative strength are responsible for the different response peaks in the left panel of figure 11.

RWDDM also reproduces the superposition observed when FI response curves are normalized by maximum response rate and time to reinforcement (top right panel of figure 11).

#### 3.7.2 Discussion

Gibbon and Balsam (1981) attributed the ISI effect to the expectancy to reinforcement. A specific reinforcer carries, according to their view, an amount of expectancy *H*. This expectancy is spread back in time over the stimulus that signals US occurrence. Hence, for a CS of fixed duration *T* and US with expectancy amount *H*, the total expectancy during the CS is *h_T_* = *H/T*. Our RWDDM account follows the same principles. The time to reinforcement *T* is computed by the ratio between the accumulation height at time of reinforcement Ψ(*t*^∗^) and the timer slope at the current trial *A(n)*. This leads to the asymptote of learning in equation 9 being set to *HA_i_*(*n*)/Ψ_*i*_(*t*^∗^). Superimposition of the response curves follows directly in RWDDM from the nature of noise in the linear accumulator. This noise guarantees that the time estimate produced by the model is timescale invariant (Simen et al., 2013).

The ISI effect can also be explained by the TD model with the Presence representation (Sutton and Barto, 1990) and with the more recently developed Microstimuli representation (Ludvig et al., 2012). The Presence representation consists of a single element *x* which has the value 1 when the CS is present, and 0 otherwise. Its associative strength *V* is updated by the TD rule at every time step within a trial. In longer trials (longer FIs) the strength *V* will decay more, since it is updated more times in the absence of the US. This will lead to a lower asymptotic value for *V*. However, Presence TD cannot account for the superimposition of intratrial response curves. The CSC-TD fares even worse, unable to account for either ISI effect or superimposition (see Ludvig et al., 2012, for a comparison between MS, CSC and Presence TD). The Microstimuli representation treats the stimulus as if it were composed of many units activated in sequence. Their activations follow a Gaussian shape which partially overlap. Later units have lower peaks and are wider than earlier ones. Because the number of Microstimuli are fixed, in longer FIs there is less temporal resolution which causes the US prediction to be lower than in shorter FIs, so it can explain the ISI effect. MS-TD’s account of superimposition is only partial, although clearly better than CSC and Presence-TD.

LeT in its current version lacks a mechanism to produce decreasing response peaks with increasing FIs. But it can account very well for superimposition, since its time representation is timescale invariant. The earlier version of Modular Theory, called Packet Theory, has been shown to produce the ISI effect (see top row of figure 3 in Kirkpatrick, 2002). This prediction comes from longer interval durations decreasing the probability of response packet generation in the model. MoT is also timescale invariant, so it generates superimposition quite easily.

To summarise, the ISI effect is explained either by time setting the asymptote of learning (RWDDM) or by a time representation that gets more diffuse with time, lowering the US prediction (MS-TD). Superimposition is explained either by the type of noise in the linear accumulator (RWDDM, LeT) or by stimulus units which have an approximately timescale invariant activation profile (MS-TD).

### 3.8 Mixed FI

Procedures where a stimulus signals reinforcement at more than one location in time are called mixed FI or two-valued interval schedules. A mixed FI involves only one CS which could be of short or long duration, and the subject has no way of knowing which duration it is currently experiencing until the US is delivered. Catania and Reynolds (1968) conditioned pigeons in a mixed FI and reported a pattern of responding during the long CS that resembles a combination of two distinct FIs (with two peaks) when the separation between the intervals was in the ratio 8:1 but not at smaller proportions. Cheng et al. (1993) found a similar result (experiment 2) when the intervals were in 5:1 proportion and Leak and Gibbon (1995) showed that with intervals in the 8:1 proportion the scalar property (measured by the CV) holds approximately even for three-valued interval schedules. Whitaker et al. (2003) ran three experiments with Mixed FIs in rats and found two peaks with the same CV when the proportion between the durations was greater than 4:1, but not for smaller proportions. They also found that the peak height at the short duration was higher than at the long duration in most cases. Whitaker et al. (2008) used intervals in the very small proportion 2:1 and still found two peaks that became more distinct when the short interval was presented more often than the long.

These results are interesting because they challenge in particular models of timing. They have served to provide evidence in favour of SET, and against BeT and the first version of LeT (Leak and Gibbon, 1995). Subsequently, they provided motivation for the development of the current version of LeT Machado et al. (2009). LeT can now account for the multiple response peaks in Mixed FIs, and their superimposition, but it cannot produce peaks with decreasing heights. Modular Theory has the necessary mechanisms to account for all the features of the data above. The TD models, MS and CSC, could both account for multiple peaks, but their account of superimposition would vary, with MS being superior than CSC. We show next that RWDDM can account for all features of the data in Mixed FIs.

#### 3.8.1 Simulations

In this simulation one CS was used which was followed by reinforcement either after 15 or 75 seconds randomly chosen, a proportion of 5:1. Our assumption was that in Mixed FI experiments subjects form two independent stimulus representations, one for the short interval *x_S_*, and another for the long interval *x_L_*, each with its respective associative strength (*V_S_*, *V_L_*) and timer (Ψ_*S*_, Ψ_*L*_). At CS onset, both timers begin timing, generating the two representations *x_S_* and *x_L_*, and at each point in time behaviour is guided by the representation with the highest activation value. When a reinforcement occurs, the CS representation with the highest activation value is the one to which credit is assigned.

The left panel of figure 12 shows the simulated responses averaged over 50 trials of the long 75-second duration. Two peaks, centred roughly at 15 and 75 seconds, of decreasing heights and increasing widths are clearly seen, matching roughly with the data (right panel).

**Figure 12:**
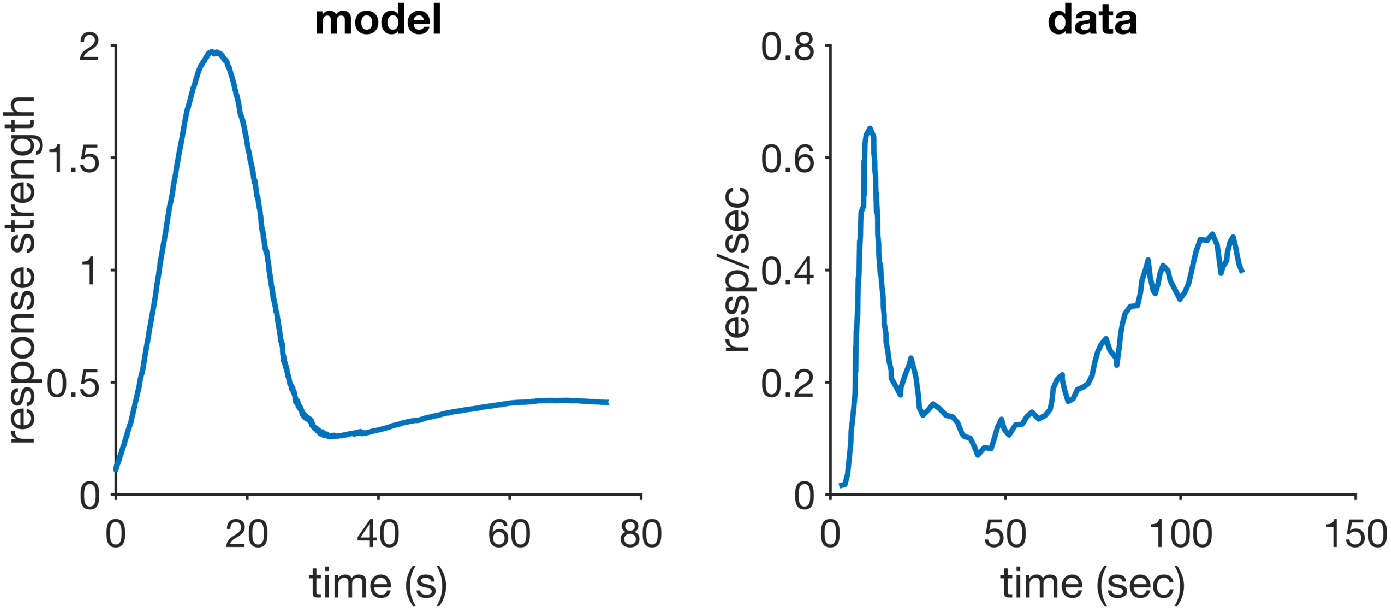
Mixed FI. Left: simulated response strength during long trials. Right: response strength data from a mixed FI experiment, redrawn from figure 3 in Leak and Gibbon (1995). Model parameters: *α_t_* = 0.2, *α_V_* = 0.1, *μ* = 1, **σ** = 0. 425, *m* = 0. 2, *H* = 30.

#### 3.8.2 Discussion

RWDDM’s mechanism for dealing with mixed FIs is in essence the same as for single FIs. The only difference is that instead of only one timer (and CS representation) in Mixed FIs RWDDM uses as many timers (and CS representations) as rewards. We have not however addressed explicitly how one CS can give rise to two distinct representations. One possible explanation is that the slope adaptation rule (equation (6)) is only applied when the difference between the two intervals is below a certain amount. If the difference is above this amount, then the model would create a new representation. In fact, the data reviewed here suggests that animals may not be able to distinguish two intervals if they are in proportion below 2:1.

To the best of our knowledge, the only other model from our analysis set that has tried to address the behaviour in mixed FIs is LeT. Machado et al. (2009) have succeeded in obtaining the two peaks with the same CV using LeT. Their account relies on a single accumulator in the form of a series of states activated at a fixed rate. This rate is fixed within a trial, but varies from trial to trial. After repeated training with a mixed FI, the states around the reinforced times receive on average more associative strength than the ones away from them. This activation pattern generates the response peaks seen in the data. However, as the authors note, ‘in mixed-FI schedules, the response rate [produced by LeT] at the first peak is equal to or lower than the response rate at the second peak, but never higher,’ which is the opposite of what the data shows. The authors suggest that a decaying arousal function might need to be added to the model so as to allow response rate to decay with interval duration.

Modular Theory is capable of accounting for the behaviour in Mixed FIs since its pattern memory for time is based on SET, which has been shown to account for these data (Leak and Gibbon, 1995). MoT’s account is similar to RWDDM’s in that both rely on a separate accumulator (and memory) for each time of reinforcement. CSC-TD would likely produce two peaks, since it relies on a perfect discretization of time into as many units as time-steps. But the curves would not superimpose when scaled as there is no mechanism to account for timescale invariance. MS-TD would also account for the two peaks but superimposition would likely not be fully obtained as its simulations of the ISI effect have only partially reproduced it (see section 3.7 and Ludvig et al., 2012).

### 3.9 VI and FI

Schedules of reinforcement specify the conditions of reinforcement delivery. There are a number of different types of schedules, some are based on the time elapsed between reinforcements, some on the number of responses emitted between reinforcements, but there can be other possibilities. Of particular interest for a timing and conditioning model are the two most commonly used time-based schedules: variable and fixed interval. Variable Interval schedules of reinforcement (VI) consist in the delivery of a US following a CS that varies in duration from trial to trial. The CS durations are usually derived from an arithmetic or geometric sequence. In contrast, Fixed Interval schedules of reinforcement (FI) use a CS of fixed duration in all trials. Skinner and Ferster (2015) reported that VIs tend to produce behaviour with a constant rate throughout the trial, whilst FIs produce scalloped curves with a pause following each reinforcement and a rapid increase in rate until the next reinforcement.

Catania and Reynolds (1968) performed a detailed analysis of behaviour under VIs and found that response rate declined with the average reinforcement rate. Within a trial response frequency increased with time, following approximately a negatively accelerated curve. When normalized by maximum response rate and time to reinforcement, these curves showed a considerable degree of superimposition.

Matell et al. (2014) trained rats on a VI in which intervals were sampled from an uniform distribution *U* (15, 45), and then tested using a peak procedure. They compared the VI response peak curve to the peak curve from a control group trained on an FI 30 (the mean of the VI distribution). Although the two curves were not significantly different statistically, the VI response peak curve peaked slightly earlier and was slightly higher than the control group.

Jennings et al. (2013) compared timing performance between VI and FI in three experiments, but found VI timing only in a VI where the average interval was 30 seconds. The other experiments from the same paper produced results more in agreement with the earlier work by Skinner and Ferster (2015) showing a constant rate of responding during VI trials.

Taken together, these studies appear to show that timing may sometimes be present during VI schedules. In this case, animals appear to be learning the average of the interval distribution. Here we demonstrate with simulations that

RWDDM can account for such findings. The only other model in our analysis set that can account for this result is Modular Theory.

#### 3.9.1 Simulations

In this simulation a random VI was produced by sampling intervals from a discrete uniform distribution U(15, 45). Non-reinforced peak trials of duration 135 seconds were interspersed during the VI, with a probability of 0.25. Our assumption here is that subjects will keep adapting the timer rate *A* over trials. In this case, equation (6) calculates the exponential moving harmonic average of the CS durations. Since it is a moving average, the predicted peak time will depend on the actual intervals used and their presentation order, but the nonmoving harmonic average of all intervals is 27.1 seconds. This is earlier than the arithmetic average (30 seconds), which is in line with the trend observed in the data by Matell et al. (2014).

Figure 13 (top left panel) compares the response strength averaged over peak trials in the VI and in a regular peak procedure with FI 30. The VI peak is higher and slightly earlier (at roughly 29.68 sec) than the FI peak, matching roughly with the data (bottom row). When normalized both by peak height and time the curves show the superimposition (top right panel) also seen in the data.

**Figure 13:**
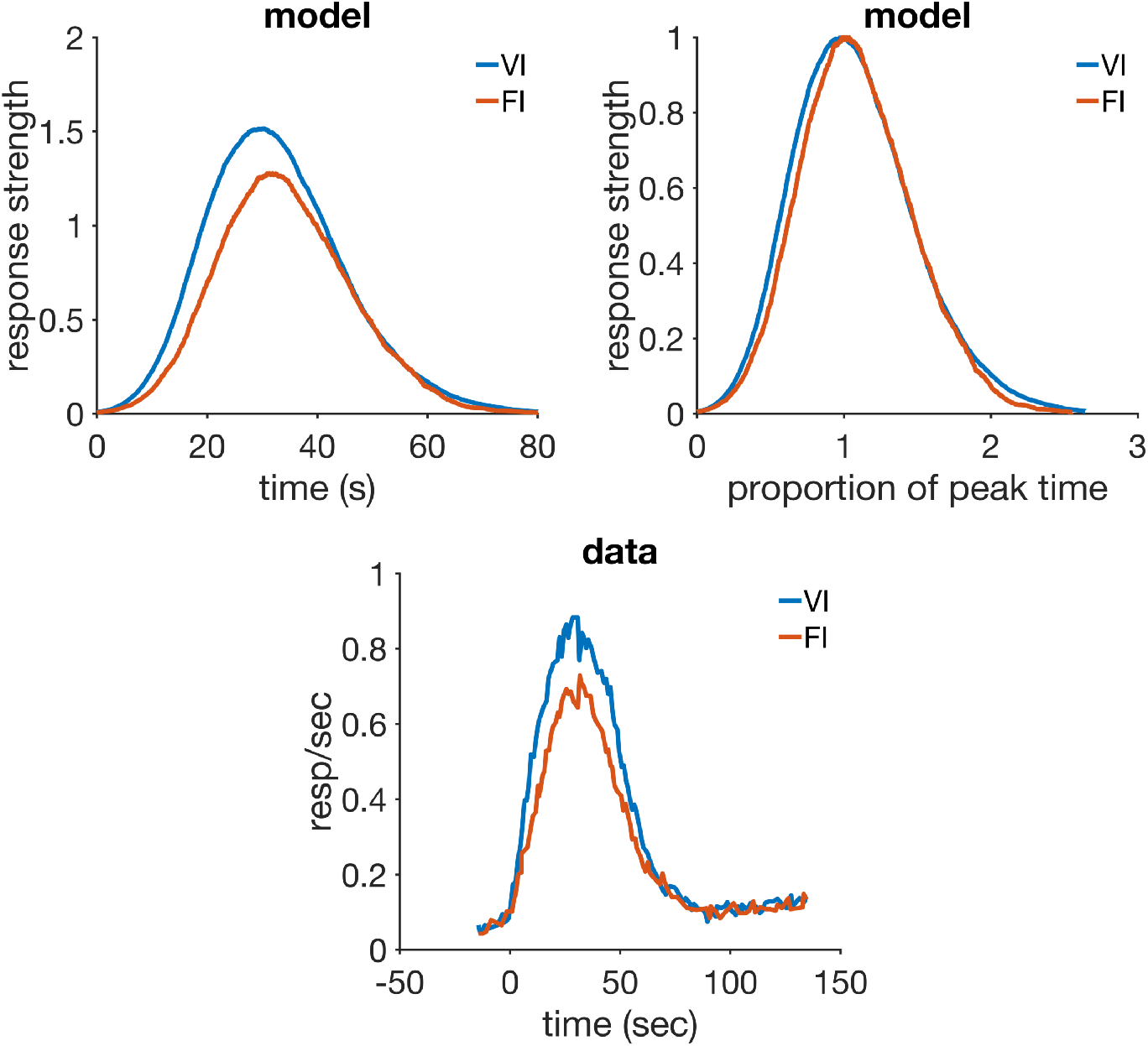
VI and FI. Top row: simulated average response strength during peak trials (left), and the same data plotted after both axes are normalized (right). Bottom row: average response strength data from an experiment in VI and FI, redrawn from figure 1 in Matell et al. (2014). Model parameters: *α_t_* = 0.1, *α_V_* = 0.1, **μ** = 1, **σ** = 0.3, *m* = 0.2, *H* = 40.

#### 3.9.2 Discussion

The model predicts a harmonic mean value for the position of the response peak, which is always less than the arithmetic mean, but because it is a weighted moving average the actual value may vary. As we saw in the simulations, the VI response curve peaked at a value (29.68 sec) very near the arithmetic mean of the intervals (30 sec). This may explain the trend observed in the data by Matell et al. (2014). However, because that trend was not statistically significant, further experiments would be needed to establish if the response peak during VIs is nearer to the harmonic or the arithmetic mean.

Taken together, these results are more easily accommodated by theories that can store an average of CS durations like RWDDM. Modular Theory is such an example, since it also stores an average of intervals in its pattern memory. Other models such as LeT and MS or CSC-TD would struggle with this result. The CS representation in these models break down the CS into a sequence of units activated serially in time. With a uniform distribution of CS durations associative strength would likely be spread broadly over the weights that cover the interval, generating a broader pattern of responses that would not be centred on the mean.

### 3.10 Temporal Averaging

Although animals are able to time different durations simultaneously, as seen in mixed FIs, paradoxically under certain circumstances a type of temporal averaging can be observed. This is a relatively new and important phenomenon, which challenges in particular theories of timing to propose a mechanism that can explain such averaging.

When rats are trained using two distinct stimulus modalities, a visual stimulus (a light) and an auditory (a tone), each signalling reinforcement at a different time, responding during compound presentations of both stimuli peaks roughly in the middle of both durations (Swanton et al., 2009). This intermediate response curve to the compound superimposes with the two other single stimulus curves when normalized, suggesting that the animal is timing only one average duration. The type of average being computed appears to be modulated by the reinforcement probabilities associated with each stimulus duration, with the weighted geometric average fitting the data better than a weighted arithmetic average or a non-weighted average (Swanton and Matell, 2011; Matell and Henning, 2013; Matell and Kurti, 2014). Significantly, temporal averaging in rats is only consistently observed when the auditory stimulus signals the short interval and the visual stimulus signals the long interval (Swanton and Matell, 2011; Delamater and Nicolas, 2015). Even when each stimulus is associated with a different response option (light reinforced with a left nosepoke, tone with a right) rats still tend to mix the temporal information during compound trials (De Corte and Matell, 2016).

We do not make a strong claim about RWDDM’s ability to explain this data. Rather, we show that it has the necessary elements from which an account can begin to be formulated. MoT also has similar elements from which an account can be built. CSC-TD, MS-TD and LeT do not appear to be equipped to deal with this phenomenon.

#### 3.10.1 Simulations

In RWDDM the accumulator is the mechanism that marks the passage of time. The temporal proximity to an event is determined by how close the level of accumulation is to a fixed threshold value. A CS that signals reward later than another CS, will have a lower rate (*A*_low_) of accumulation than the shorter CS (*A*_high_). Because in RWDDM associative strength is set by time to reward, the two CSs will also have different associative strengths, *V*_low_ and *V*_high_ respectively. We may assume that under temporal averaging circumstances the stimuli are of such nature that they cause the subject to integrate their information. At the start of the compound trials, the ambiguity presented by the compound stimulus may cause the representations of the two component stimuli to be only partially retrieved. If the subject fails to represent the two stimuli separately, the result may be the formation of a single representation composed by only a fraction of the timing rate *A* and associative strength *V* of each individual stimulus. The fractions are then added into one single rate and one single associative strength, and processed as if they were the components of a single stimulus representation. For the simulation below, we assume that the fractions added are exactly half of their individual values: *A*_compound_ = *A*_low_/2 + *A*_high_/2, and *V*_compound_ = *V*_low_/2 + *V*_high_/2.

We used a long CS of duration 20 seconds and a short CS of duration 10. We simulated a peak procedure with each CS and with the compound. A plot of the response strength averaged over peak trials is shown in the top left panel of figure 14. The three peaks scale when normalized (top right panel).

**Figure 14:**
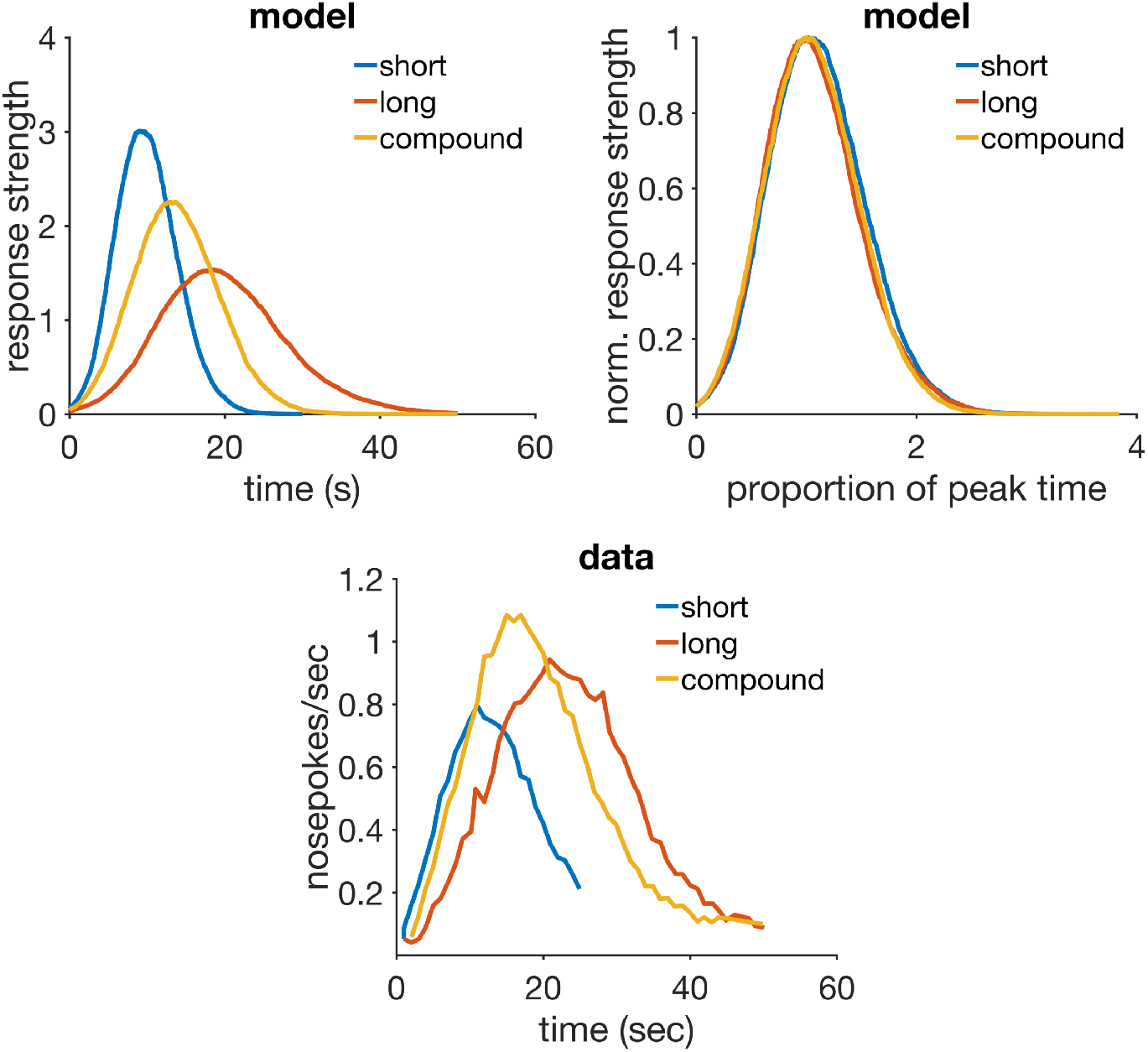
Temporal averaging. Top row: simulated response strength averaged over peak trials in temporal averaging (left), and the same data normalized by maximum response strength and peak time (right). Bottom row: peak trial response strength data from an experiment in temporal averaging, redrawn from figure 1 in Swanton et al. (2009). Model parameters: *α_t_* = 0.2, *α_V_* = 0.1, **μ** = 1, **σ** = 0. 35, *m* = 0. 2, *H* = 30.

The peak of the compound is roughly at 13.33 sec, which would be the expected value for an averaged rate *A* = (1/10 + 1/20)/2, the harmonic average of the intervals. The height of the compound peak is also at an intermediate level between the two end peaks. The simulations match roughly with the data (bottom row of figure 14)

#### 3.10.2 Discussion

The assumption we made here, that temporal averaging is the result of only one accumulator being active during the compounds and fed with half the rate for each of the stimuli, is plausible and can accommodate the main features of the data. However, given the evidence from mixed FIs it seems animals are capable of keeping multiple timers running in parallel, without averaging their rates. Also, if averaging of rates always happened during compounds, then the explanation provided by RWDDM for the left shift in the response curve in the compound peak procedure would not hold. We suggest one possible way of interpreting these three phenomena based on a failure of representation selection caused by the ambiguity of the signal. In mixed FIs there is one single CS that signals two rewards at very different times. There is not much ambiguity in how to interpret the signal, so the subject keeps two timers running in parallel. In the case of compounds formed by individual CSs that signal reward at the same time, as in the compound peak procedure, there is also not much ambiguity. There’s very little difference between the time memories evoked by the CSs, so choosing only one, the faster one, leaves no ambiguity as to which CS is signalling reward. In the case of compounds formed by individual CSs of different modalities that signal reward at different times, the ambiguity might be such that cannot be resolved easily. The information from each CS may then be only partially retrieved and added into one representation, resulting in temporal averaging.

As mentioned previously, this is not a strong account of the conditions that generate temporal averaging. But whatever the final word on this may be, RWDDM has components that allow it to generate averaging and timescale invariance. However, RWDDM predicts this average to be the harmonic mean, and not the geometric mean weighted by reinforcement probabilities that has been frequently found (Swanton and Matell, 2011; Matell and Henning, 2013; Matell and Kurti, 2014). Also, Matell and Henning (2013) reported evidence of summation of response rates during the compound trials. In our simulations here we assumed that equal fractions were taken of the rates of each CS, resulting in a combined non-weighted harmonic average of rates, but different fractions (or weights) may be taken. In particular, the data indicates that the weights are set by the reinforcement probabilities of each individual stimulus. Since this information is stored in the associative strength *V*, we could assume the subject integrates the two timer rates as follows:

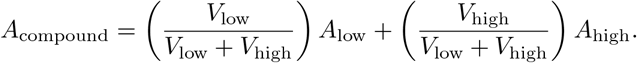

Although this would produce a weighted average, it is still a weighted harmonic average of the intervals and not a weighted geometric average found in the data, so the account given by RWDDM would still be partial. As for the summation of response rates observed in the compound trials, this could be explained by RWDDM if instead of taking a fraction of the *V* values for each stimulus to form the *V* compound, the subject simply summed, or partially summed, both *V* values.

Another model that is equipped to deal with averaging is Modular Theory. If we allow for one single accumulator fed by one half of each time memory, then MoT would predict a peak of responding at the arithmetic mean of the two intervals. A weighted average could also be obtained following the procedure we sketched above for RWDDM. However, this would yield a weighted arithmetic mean, and not the weighted geometric mean obtained in the data. As for timescale invariance, MoT relies on a noisy timer threshold whose mean is always a fixed proportion of the time memory, with a standard deviation proportional to this mean. Therefore, timescale invariance is guaranteed for all time memories, averaged or not.

LeT would not be able to explain temporal averaging without modifications. It cannot change its average transition rate between states without compromising timescale invariance. Without changing the transition rate it is difficult to see how else LeT could account for a different timing in the presence of the compound. CSC-TD and MS-TD also lack any mechanism that could be used to account for temporal averaging.

### 3.11 Summary of Results and Analysis

Table 4 summarizes the results from the simulations. RWDDM was able to reproduce the main features of the data in 8 out of the 10 experiments. In the other 2 the model was able to partially account for the data.

**Table 4:** Summary of main simulation results and comparison with other models. Notes: (1) if learning rate is allowed to vary.

| phenomenon | RWDDM | CSC-TD | MS-TD | LeT | MoT | explaining mechanism |
| --- | --- | --- | --- | --- | --- | --- |
| faster reac-<br>quisition | yes | yes<sup>1</sup> | yes<sup>1</sup> | yes<sup>1</sup> | yes<sup>1</sup> | time-adaptive stimu-<br>lus representation or<br>changes in learning<br>rate |
| time change<br>in extinction | yes | no | no | no | yes | separate rules for time<br>adaptation and asso-<br>ciative strength |
| latent inhibi-<br>tion and tim-<br>ing | part. | no | no | no | no | PH rule and separate<br>rules for time adap-<br>tation and associative<br>strength |
| blocking with<br>diff. dura-<br>tions | part. | yes | yes | no | no | RW rule and ability to<br>time any stimulus or<br>distributed time rep-<br>resentation |
| time spec. of<br>conditioned<br>inhibition | yes | yes | yes | no | no | RW rule and con-<br>centrated memory for<br>time or distributed<br>time representation |
| compound<br>peak proce-<br>dure | yes | no | no | yes | yes | intertrial variability in<br>time estimation |
| ISI effect and<br>superimposi-<br>tion | yes | no | part. | part. | yes | asymptote of assoc.<br>strength set by time<br>and accumulator noise<br>or time representation<br>that gets diffuse with<br>longer time |
| mixed FI | yes | part. | part. | part. | yes | ability to generate<br>multiple time rep-<br>resentations or a<br>single distributed<br>time representation |
| VI and FI | yes | no | no | no | yes | memory that stores<br>average of intervals |
| temporal av-<br>eraging | yes | no | no | no | yes | memory that stores<br>average of intervals<br>and the accumulator |

To allow for comparison we have offered qualitative predictions for the other 4 models in table 4. It is important to note that for most of the 10 phenomena analysed here simulations using these models are not available in the literature. Although we have tried our best to provide predictions based on our understanding of these models, we have not actually simulated them. Therefore it is possible that in some cases a model may produce results that we did not foresee if the right set of parameters is found or some of the assumptions are relaxed. It is also possible that some simple modifications might allow the models to explain the data. We endeavoured to point out some such modifications that seem likely to work when discussing the simulation results above, but we do not make predictions based on them because the purpose here is only to provide a comparison of the current mechanisms of each model and therefore encourage future work on model improvement. With that in mind, Modular Theory has fared best after RWDDM, being able to account for 7 out of the 10 experiments. MS-TD and CSC-TD shared the second place with 3 out of 10. LeT came in last, able to account for 2 experiments. The last column of table 4 identifies the main mechanisms responsible for successfully accounting for each phenomenon.

## 4 General Discussion

RWDDM was able to reproduce faster reacquisition due to its memory for time being conserved during extinction. This memory is used to activate the stimulus representation. Learning is slower in acquisition because RWDDM increases the activation in the stimulus representation gradually over the trials. The stimulus representation needs to be ‘built up’ first, and this process depends on learning the timing of the US. Extinction eliminates associative strength but leaves the time memory, hence the stimulus representation, intact. Reacquisition proceeds faster because the stimulus representation does not need to be built up again. Other models explain this by allowing the associative strength learning rate to be faster in reacquisition.

Time change in extinction was accounted for because of RWDDM’s ability to time CS duration independently from US associations. Time is learned entirely by time markers. The TD models and LeT do not make this separation. These models do not have a mechanism to time stimuli without the US stamping in the changes.

Improved timing in latent inhibition was also accounted by RWDDM’s ability to learn timing independently of associations. Preexposure allows the model to build its time representation, which is later expressed by behaviour during the acquisition phase. The only other model that learns to time independently of associations is MoT, but it does not have a mechanism to explain the latent inhibition effect. The latent inhibition effect alone, i.e. the initial decrement in the acquisition curve of a preexposed stimulus, was made possible in RWDDM by using the P-H rule to change the learning rate for associative strength. The use of the P-H rule instead of the RW would certainly have other theoretical implications for the general theory we are introducing in this paper, but we have used it only in this case. We will make further comments in the conclusion. Blocking with different durations was easily accounted in one condition, the short blocked and long blocking CS. The blocking effect in this condition followed from the summation term in the RW rule. For the other condition, long blocked and short blocking CS, a straight application of the model did not yield the results expected. But the experimental results leave open the possibility that this might be a case of second-order conditioning, where the summation term in RW does not play a role. In this case, RWDDM is well placed to explain the results, since it can time the whole sequence of stimuli. The only other models capable of explaining these results were the TD models.

The time specificity in conditioned inhibition was very well accounted for by the combination of the summation term in the RW rule, which allowed for inhibition to develop, and the independent timing mechanism in RWDDM that allowed it to time US omission. However, the alternative account provided by the different time representation in the TD models was also successful. The other theories failed here for the same reason as in blocking, they lack a rule like RW that can deal with compound stimuli effects.

The response curves centred at the mean of intervals in the VI procedure was well accounted by the ability of RWDDM to learn the average of intervals. This ability is only present in Modular Theory, making it the only other model able to account for the results here.

In the case of temporal averaging, RWDDM was able to account for the general features of the phenomenon, namely a response curve that peaks at the average of the intervals signalled by the compound stimulus. However, RWDDM predicts the peak to be at the harmonic mean, whilst some experimental results suggest it happens at the geometric mean. RWDDM’s account of temporal averaging was hypothesised as the result of ambiguity in the signal. In trying to resolve whether the compound should be treated as a single stimulus or as two separate stimuli, the subject settles on using one accumulator that is fed partial timing information from both stimuli. Other hypothesis might turn out to be more adequate, but this is one possibility that fits well with the RWDDM framework. The only other model that would produce averaging under the same hypothesis is MoT.

The classic ISI effect followed from two mechanisms in RWDDM. The lower response curves during longer stimuli was explained by time setting the asymptote of associative learning. The larger spread of response curves during longer stimuli and the superimposition of normalised curves follows from RWDDM’s timescale invariant time representation. The noise in RWDDM’s accumulator decreases with the interval being timed, in such a way that it results in timescale invariance of the response curves. Modular Theory can also reproduce all features in the data. This is because it relies on a timescale invariant response rule function that generates less responding in longer intervals. LeT can account for superimposition, but it does not have a mechanism to account for the lower curves in longer stimuli. MS-TD can account for both elements because of the form of its microstimuli representation.

The double peaks observed in the response curves during mixed FIs is explained by RWDDM using simultaneous timing. It generates two different representations, one for each reward. Thus, it can account for mixed FIs by the same principles used to account for the ISI effect and simple FI schedules. Modular Theory takes the same approach of simultaneous timing and is also successful. The TD models and LeT can provide a partial account due to their distributed time representation. But timescale invariance of the peaks is not observed in CSC-TD and only approximately in MS-TD. LeT produces the timescale invariance but not the decrease in peak height with time.

The left shift of response curves seen in compound peak procedure and dis-inhibition of delay was well accounted for by RWDDM. It did so because of intertrial variability in noise estimation. By choosing in every compound trial the time memory that predicts reward sooner, RWDDM produces the left shift in response. The only other models that can appeal to the same principle to explain it are LeT and MoT.

The superiority of RWDDM and MoT in explaining the majority of the phenomena analysed highlights the importance of some of their shared mechanisms. Both models have separate rules for updating time and associative strength. This makes them capable of timing any stimuli, independent of changes in associative strength. Both models represent psychological time as linearly related to physical time through the theoretical construct of the accumulator. Their memory for time stores a moving average of the experienced intervals. They both allow for intertrial variability in time estimation. Among their differences, only one proved crucial in discriminating the two models in the experiments analysed here: the lack of a mechanism in MoT to account for stimulus compounds. RWDDM uses the RW rule, which was developed to deal with phenomena such as blocking and inhibition, whilst MoT uses the linear operator, a historically earlier association rule that cannot handle compounds. This was the single difference that caused the difference between MoT and RWDDM in number of phenomena explained.

MS-TD came in third place in number of phenomena successfully explained, but the gap between it and MoT was comparatively high, with MoT being almost twice more successful than MS-TD. CSC-TD came just half a point below MS-TD. This is certainly a result of their similarities. The only difference between these two TD models is in their time representation. However, this different representation allowed MS-TD to explain only one more phenomenon than CSC-TD, the ISI effect. Therefore, in the set of experiments analysed here MS-TD did not show a significant improvement on CSC-TD. This does not mean that MS-TD is not a significant improvement on CSC-TD overall. Its superior account of timing is significant. But the set of experiments chosen here are particularly challenging even for a dedicated timing theory, so they raise the bar even higher. The strength of the TD models was in accounting for compound phenomena of blocking and inhibition, due to their RW rule for association. Their weaknesses was that they rely on changes in associative strength to express changes in timing. This prevented them from explaining time change in extinction and improved timing in latent inhibition. They both lack a memory to store the average of intervals, so they could not explain behaviour in VI schedules. Finally, their lack of trial to trial variability in time estimation prevented them from accounting for the left-shift in the compound peak procedure.

With respect to the number of successes only, LeT came in last. The results allowed us to identify at least four limitations in LeT’s current formulation. The first is that it ties its time representation to changes in associative strength. This prevented it to explain time change in extinction and improved timing in latent inhibition. The second limitation is that it relies on the linear operator rule for associative strength, which prevented it from accounting for blocking and time specificity in conditioned inhibition. Thirdly, its distributed memory for time does not store the average of the intervals seen. This prevented it from accounting for the behaviour in VI. Lastly, it doesn’t have a mechanism to explain the decrease in peak height of the response curves with longer ISIs. However, as a timing model, LeT’s strength is in explaining timescale invariance. If it can be made to overcome at least the weakness of its associative learning rule, for example by also adopting the RW to update associative strength, LeT could be on a par with the TD models.

RWDDM faced a couple of difficulties in explaining the set of phenomena analysed here. In latent inhibition the model was able to learn the timing for the preexposed CS, but our choice of CS representation translates this into a response curve that does not fully match the data. A better solution might involve a two-state CS representation, one state for the early stages of training and the other for the latter stages. Finally, RWDDM could not account for the lack of blocking with a long blocked CS and a short blocking CS. One possible solution that does not require changing the model is to treat the blocking CS as a secondary reinforcer.

One relevant phenomenon that we did not explore here is the peak procedure. In particular, Balci et al. (2009) have produced evidence that in the long peak trials animals don’t stop responding immediately after the expected reward time, but instead take a number of peak trials to learn to stop. The Gaussian function *x_i_* (Ψ_*i*_) used as the CS representation in RWDDM ensures that CR levels will begin to decrease after Ψ*_i_* (*t*) crosses threshold *θ* without any learning. To address the findings in Balci et al. (2009) the RWDDM CS representation could be changed to a sigmoid, saturating after the timer Ψ(*t*) crosses a first threshold. A second threshold could then be introduced to mark the time to stop responding. When the timer crosses this stop threshold the saturation process in the CS representation would stop and a decay process would begin. This however would still be an incomplete account, as a mechanism would be needed to explain the learning of the second threshold. But if such a CS representation was used, the model would also fit a larger body of data coming from studies that analyse responding during individual trials of the peak procedure. Schneider (1969) and subsequently Gibbon and Church (1990) and others (Cheng and Westwood, 1993; Matell et al., 2006) have argued that the pattern of responding is better characterized not by a Gaussian but instead by an approximate square-wave function, with a low-high-low response frequency pattern. It can be shown that by introducing a stop threshold to the timer Ψ*_i_* (*t*), the TDDM timer (used in RWDDM) can fit the data on times of start and stop responding (Luzardo et al., 2017). Alternatively, the accumulator Ψ*_i_* (*t*) itself could be used as the CS representation, replacing *x_i_* in equations (9) and (10). In this case, an upper absorbing boundary would need to be set on the accumulator to prevent response strength increasing considerably in the first few trials following a CS duration increase for example. Also, such a choice of CS representation would cause within-trial responding to become linear, rather than the more commonly observed sigmoidal pattern. If a sigmoidal response curve is to be preserved, a different choice of response function would be required.

Another phenomenon that we did not address but deserves mention is the timescale invariance of the acquisition process (Gallistel and Gibbon, 2000). It refers to the general finding that the number of trials required until an acquisition criterion is met depends on the ratio of intertrial (or context) and trial durations, the I/T ratio (Gibbon, 1977; Lattal, 1999; Holland, 2000). Gibbon and Balsam (1981) provided an account for this that postulates a decision process based on the reward expectancy signalled by the stimulus versus the one signalled by the context. A ratio between the two expectancies is calculated, and once the ratio exceeds a certain value, acquisition starts. If the same postulate of a decision ratio of reward expectancies is made, RWDDM may account for the I/T ratio in a similar manner. If we assume that animals time the interval between USs (the context or I duration) with rate *A_I_* (_*n*_) and also the CS duration as usual with rate *A_T_* (*n*), then we can form the ratio *r(n) = A_*T*_(n)/A_*I*_(n)*. As the number of trials *n* increase, the *A* rates converge to their asymptotic values, and the ratio *r* will converge to *A_T_/A_I_* = (1/_*T*_)/(1/_*I*_) = *I/T*. This is essentially the same account given by Gibbon and Balsam (1981), with the timer rates *A_T_* and *A_I_* substituting Gibbon and Balsam’s expectancies *H/T* and *H/C*.

At least three testable RWDDM predictions came out from the simulations reported here. The first concerns blocking with different durations. A long blocked CS will not be blocked by a short co-terminating blocking CS, and two peaks in responding will be observed during test trials with the blocked CS: one at the time the short blocking CS would normally start, and another at the end of the blocked CS. The second prediction is that conditioned inhibition is the exact opposite of excitation. This means that the behaviour produced by inhibition is timed in the same manner as in excitation. Finally, in temporal averaging the response peak in the compound stimulus should be at the harmonic average, or weighted harmonic average.

RWDDM is, to the best of our knowledge, the first time the RW associative learning rule is coupled with a accumulator-based timing theory. An important implication of this effort for associative learning is that it allows for a richer analysis of the effects of timing in compound stimuli experiments. Here we have analysed blocking and conditioned inhibition, but there is evidence suggesting time may have important effects in other cue-competition phenomena such as overshadowing (Kehoe and James, 1983; Jennings et al., 2007). Timing effects in compounds has until now received somewhat little attention, with many published experimental studies reporting only aggregate response measures. This is perhaps to be expected, since most associative learning models that can handle compounds do not have any, or a rich enough, time representation. RWDDM is an attempt at filling this theoretical gap.

Another limitation of associative learning models is that they tend to simply postulate the timing features of the stimulus representation, without a detailed account of how these can mechanistically arise and evolve. This is the case with the CS representations of CSC-TD, MS-TD and others like C-SOP (Brandon et al., 2003). RWDDM’s adaptive timer and time-adaptive CS representation provide a fuller account of the timing mechanism and its dynamics. Another recent model that provides this level of detail is the Timing from Inverse Laplace Transform (TILT, Shankar and Howard, 2012; Howard et al., 2015). It can dynamically develop a timescale invariant representation of stimulus history using a two-layer neural network. It can also reproduce a number of important conditioning phenomena, but so far it has only been implemented with the linear operator rule for associative learning, which precludes it from accounting for cue competition phenomena.

The RWDDM architecture suggests that timing is largely independent of the process of association formation and maintenance. Associations however, according to RWDDM, depend on timing both to set the asymptote of associative strength and to build the CS representation so that it can enter into association with the US. Thus, RWDDM implies that interactions between timing and associative learning are mainly one-directional. This appears to match roughly with experimental findings. In a review Kirkpatrick (2013) found that prediction error influenced measures of time estimation only through changes in reward magnitude and devaluation, whilst effects in the other direction included the appropriate timing of CRs from start of conditioning, trial and intertrial durations affecting strength and probability of CR occurrence, and cues with different temporal information affecting cue competition.

## 5 Conclusion

In this paper we introduced a new real-time model for classical conditioning and timing. The model combines elements from two theories, the Rescorla-Wagner conditioning model and the TDDM interval timing theory.

We have simulated the model on 10 conditioning phenomena selected from the literature, which collectively represent a particular challenge for any single model to explain. The model was successful in accounting for 9, and can be made to account for the rest if simple modifications are made. The mechanisms used by other models of similar scope were evaluated to see if they could also account for the data. The only other model to reach a comparable level of success in this set of phenomena was Modular Theory. This was due to MoT and RWDDM having a significant overlap in terms of mechanisms. Our analysis suggests that certain mechanisms may also be adopted by some of the other models, thereby improving their explanatory power.

RWDDM may be improved in several ways. It is quite likely that the asymptote of learning may not be described by the simple inverse relationship to reinforcement time that we assumed. In some of the experiments modelled here, response peak seemed to decrease slower with ISI than our inverse relationship predicted. Functions other than Gaussians might be used to represent the CS, which could better fit the data in the case of latent inhibition for example. These and other theoretical issues may be better elucidated by new experiments involving compound stimuli and a manipulation of their durations, such as the experiments with blocking, compound peak procedure and temporal averaging analysed here.

We have also adopted the P-H rule in one experiment, but have not explored its application in the others. Making the P-H rule an integral part of RWDDM would add one more parameter but it would also allow RWDDM to account for other preexposure and attentional effects that the rule is designed to account. This is not a difficult modification, and we have already shown it to be feasible.

RWDDM may be regarded, like TD, as a real-time extension of RW. It adds to it the powerful timing mechanism of TDDM. But also, by making a link with a version of DDM, it shows that it may be possible to arrive at a unified account of timing, conditioning and decision making.

